# An orderly single-trial organization of population dynamics in premotor cortex predicts behavioral variability

**DOI:** 10.1101/376830

**Authors:** Ziqiang Wei, Hidehiko Inagaki, Nuo Li, Karel Svoboda, Shaul Druckmann

## Abstract

Animals are not simple input-output machines. Their responses to even very similar stimuli are variable. A key, long-standing question in neuroscience is understanding the neural correlates of such behavioral variability. To reveal these correlates, behavior and neural population must be related to one another on single trials. Such analysis is challenging due to the dynamical nature of brain function (e.g. decision making), neuronal heterogeneity and signal to noise difficulties. By analyzing population recordings from mouse frontal cortex in perceptual decision-making tasks, we show that an analysis approach tailored to the coarse grain features of the dynamics was able to reveal previously unrecognized structure in the organization of population activity. This structure was similar on error and correct trials, suggesting what may be the underlying circuit mechanisms, was able to predict multiple aspects of behavioral variability and revealed long time-scale modulation of population activity.

## Introduction

Decision making is a central behavioral paradigm in neuroscience. In such tasks, animals sample sensory inputs, generate a behavioral choice, hold it in memory, prepare an action, execute the action, and finally compare the outcome to the expected delivery of reward. These elaborate behavioral computations manifest in complex neural dynamics, occurring across multiple temporal scales and multiple brain areas^1-11^. For example, recent studies in the mouse identified anterior lateral motor (ALM) cortex, where neurons exhibit complex, heterogeneous, epoch-dependent neural dynamics, as a critical circuit node for perceptual decision tasks^9-12^. Population dynamics in ALM and connected thalamic nuclei^13^ evolve over time scales of tens to hundreds of milliseconds within a single trial. These neural dynamics reflect the confluence of sensory information, representation of upcoming movement, and other internal states. Yet due to the complexity of population dynamics and the partial sampling offered by experimental preparations, the nature of that representation and how it encodes behavioral variables and internal states remains unclear.

Across trials, identical stimuli give rise to variable neural responses and behavior. Hence, analysis that averages trials risks masking the link between behavioral and neural variability. Neural activity may explain variable behavioral outcomes, yet variability in single neuron activity, especially in neurons with low firing rates, presents a challenge to such analyses. Moreover, simultaneous recordings have consistently revealed that neural activity is diverse, both within and among brain areas and across time. This complicates efforts to relate neural dynamics to underlying computations and to behavior.

Population activity analysis methods can leverage simultaneous recordings to pool information across neurons and reveal information that is difficult to decode from raw single neuron or population activity^8,14-27^. Typically, these approaches infer intermediary variables (latent variables) from the activity of simultaneously-recorded neurons in single trials, then use these variables as the basis to predict behavioral outputs. These methods have been used to improve estimation in brain-computer interfaces^28,29^ and predict behavioral properties such as reaction time, indecision and hesitation.

In the present study, we developed a variant of latent space dynamical system methods to uncover decodable information from complex, temporally heterogenous dynamical neural recordings. We assume that population activity in single trials can be explained by the evolution in time of a lower dimensional dynamical system learned from the data that represents shared activity across neurons^30^. Our approach utilizes the conceptual and computational simplicity of linear dynamical system models, but robustly extends their efficacy by allowing the underlying dynamics to change over time. Importantly, switches between distinct dynamical systems occur at cued times, the transitions between trial epochs, when population dynamics are known to change ^9-11^ (**Fig. 1b, Fig. S1**). For instance, shortterm memory is necessary during delay epochs, but not necessarily before stimulus presentation. This approach represents a compromise between two extremes: on one hand assuming that the nature of the underlying dynamics is linear and fixed across time ^14-17,24-26^, which we found to be insufficient to capture the complexity of neural dynamics during decision-making; and on the other hand using more complex non-linear dynamics models by allowing switches between a set of different linear dynamics at any given time^18,27,31^. Models that allow switches at arbitrary times are difficult to fit, due to the necessity of tracking combinatorial number of past switching options. For instance with a set of K dynamics there could be a switch into one of K states after the first time point, and then K^2^ options after the second time point of switching from any of the first-time-point options into any of the K states, and so on.

**Figure 1.**
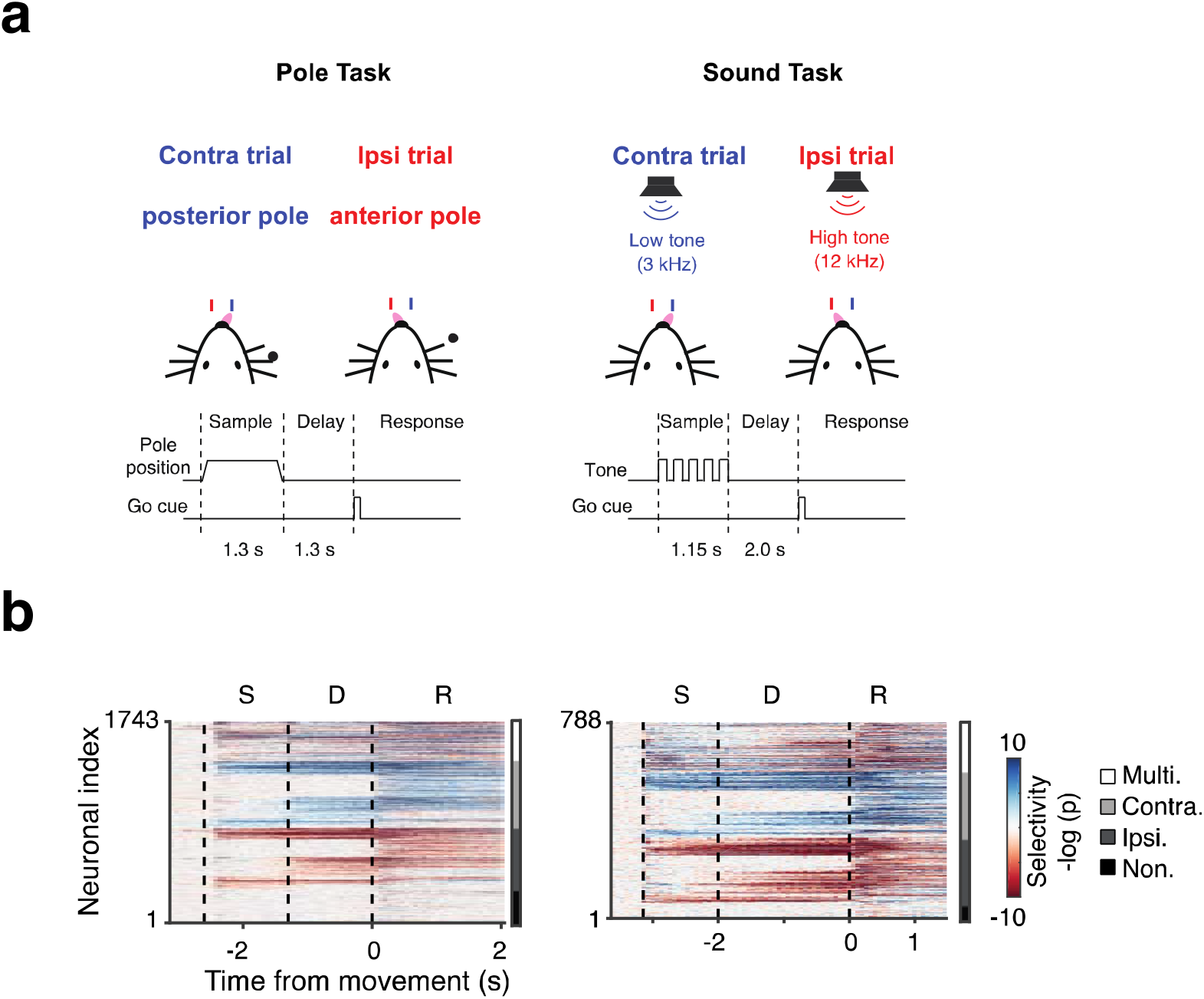
Neural activity of anterior lateral motor cortex neurons exhibits complex, heterogenous activity. **a.** Schematic description of experiments. Mice were trained to report pole position (left) or frequency of tone (right) by directional licking, following a delay period after the end of stimulus presentation (contra trial – posterior pole, lick right; ipsi trial – anterior pole, lick left). **b.** Heatmap of instantaneous trial-type selectivity, displayed as negative log of p-value, (blue: contra-preferring; red: ipsi-preferring). For visual clarity neurons were sorted by response category, shown in bar on right (black, non-selective; dark gray, ipsi-monophasic selective; light gray, contra-monophasic selective; white, multiphasic selective, where neurons show strong switches of selectivity during a task).

By analyzing simultaneous recordings from ALM, we find that a large fraction of neural variability was captured by latent space dynamical system models. The low dimensional latent variables in our model, which we refer to as the shared activity space (SAS), reveal dynamics that are robustly maintained for seconds pre- and post-decision. The dynamics in the SAS predicted multiple forms of single-trial behavioral variability, such as response latency and correctness of animal responses. Dynamics discovered during correct trials could still describe activity during error trials, and differences in the state of the SAS were large enough to predict an animal’s error hundreds of milliseconds to seconds before the actual action occurred.

Our approach revealed previously unrecognized structure within the population dynamics of ALM. The animals performed a two-alternative-forced choice task with a delay. That is, they were presented with one of two sensory stimuli during a sample period and were required to execute a specific action matched to each sensory stimulus, but only during the response period which was separated from the sample period by a delay period. To successfully perform the task, the animal need only maintain trial-type-selective dynamics from the sample to the response epochs. Nevertheless, we find that population activity maintains the rank order of individual trials across epochs even within a trial type, concurrent with maintaining trial-type selectivity. This has implications for the underlying circuit structure. First, most discrete-attractor-based models yield random dynamics around the attractor^32^. This would mix the activity levels of trials across time, which would not preserve trial order. Second, we find dynamics are consistent across long time-scales. In many sessions within-epoch dynamics repeated reliably across multiple consecutive trials, changing on timescales of many minutes. This pertains directly to the question of where and how long-timescale internal states can be found in the brain. Indeed, we inferred longer timescale behavioral measures, correct performance on a previous trial, from these dynamical models.

In summary, we find that these modes of shared population activity reveal novel properties of circuit dynamics, allow more accurate descriptions of the single-trial variability of dynamics and can be utilized to generate useful quantitative measures of the neural substrate of behavior both on short time scales in single trials and on longer time-scales.

## Results

### Complex, epoch-dependent neural dynamics in anterior lateral motor cortex

Multiple types of optogenetic manipulations during behavior have established a causal link between ALM preparatory activity and movement^9-12^. We analyzed neural recordings from mice performing one of two types of delayed discrimination tasks. In the first, mice (n = 33) reported the position of a pole (anterior or posterior) by directional licking (lick left or lick right), but they were required to respond only at the end of a delay period lasting 1.3 seconds, with an auditory ‘go’ cue signaling the onset of the response epoch. In the other task, mice (n = 6) reported the frequency of an auditory cue after a 2-second delay (**Fig. 1a**). A large proportion of ALM neurons exhibit persistent and ramping preparatory activity during the delay epoch before the movement^9-12^.

ALM neurons exhibit complex, heterogeneous dynamics. This was true for raw single-neuron or population activity, for trial-type decoding based on single-neurons, which changed substantially over time for many neurons (**Fig. 1b**), and at the level of neuronal population selectivity (**Fig. S1**). Defining neuronal selectivity as differences between the responses to trials in which the animal was instructed to lick left and trials in which the animals were instructed to lick right, we find for example that >50% of ALM neurons (n = 917/1743, pole task; n = 444/788) changed their selectivity from the delay epoch to the response epoch. At a finer temporal resolution, neurons became selective at various time points during the trial, with many changes appearing at the onset of epochs (**Fig. 1b**). At the population level, analyzing a subset of the data containing simultaneous population activity (6 - 31 units per session, 55 sessions; **Table S1)** we also found strong switches in dynamics^27^. Defining population trial-type selectivity by linear discriminant analysis (LDA) decoders at separate times, we found that decoders were stable within epochs, but diverged across epochs. These observations held true for both tasks and even when neurons were combined across sessions into a pseudo-population (**Fig. S1**).

### Neural dynamics and the shared-activity space

Such complex neural dynamics are difficult to interpret at the level of raw population activity (i.e., a set of time series representing spiking activity). We analyzed population activity through latent dynamical system models that directly model the population activity^8,14-27,31^, assuming that it can be effectively modeled as a combination of a small number of shared patterns of neural activity as described below. However, static linear dynamical system models (LDS) do not capture the complexity of ALM activity. Fully switching linear dynamical system models (sLDS, which allow switches at any time) suffer from combinatorial difficulties in fitting the model to the observed dynamics. Instead we assume static linear dynamics during each behavioral epoch, but allow these dynamics to change at cued times, in this case across behavioral epochs:

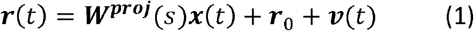

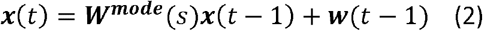

We assume that neural population activity, referred to as the full-neural space (FNS), ***r***(*t*) ∈ ***R**^N^*, can be modeled as a weighted sum of neural modes, ***x***(*t*) ∈ ***R**^M^* (*M* < *N*), i.e., a low-dimensional latent space, through a projection matrix ***W**^proj^*. We refer to this latent space of **x** as the shared activity space (SAS) since each latent mode represents common dynamics shared by many neurons. The model considers the difference between the observed activity in the FNS and the activity explained by the SAS as neural residuals, ***v***(***t***) ∈ ***R***^*N*^, that are independent for each neuron (**Fig. 2a,** top). This model can be represented as a two-layer network: The first layer, corresponding to the SAS, is represented by a set of implicit units with layer-internal dynamics, while the second layer comprises units whose activation represents the measured population activity (FNS). The activity of the second layer is the projection up of the SAS by ***W***^*proj*^ and the addition of the neural residuals (**Fig. 2a,** bottom). In SAS, temporal dynamics and interaction of latent modes are modeled as a linear dynamical system expressed by the matrix ***W***^*mode*^ and incorporating a stochastic term ***w***(***t***) ∈ ***R***^*M*^, which varies from time point to time point and across trials. This term can represent input from neurons in other parts of the brain, such as sensory areas (and is often referred to as an innovation term). Notably, both matrices that describe the dynamics ***W***^*proj*^ and ***W***^*mode*^ are epoch-dependent (indexed by s). More generally, in cases where there is no explicit epoch-design, switches can be linked to other known events, such as a shift in spatial location or a change in the environment. We refer to these models as event-dependent linear dynamical systems models (EDLDS). For the delayed response task the transitions between the epochs in each trial are the only events driving switches.

**Figure 2.**
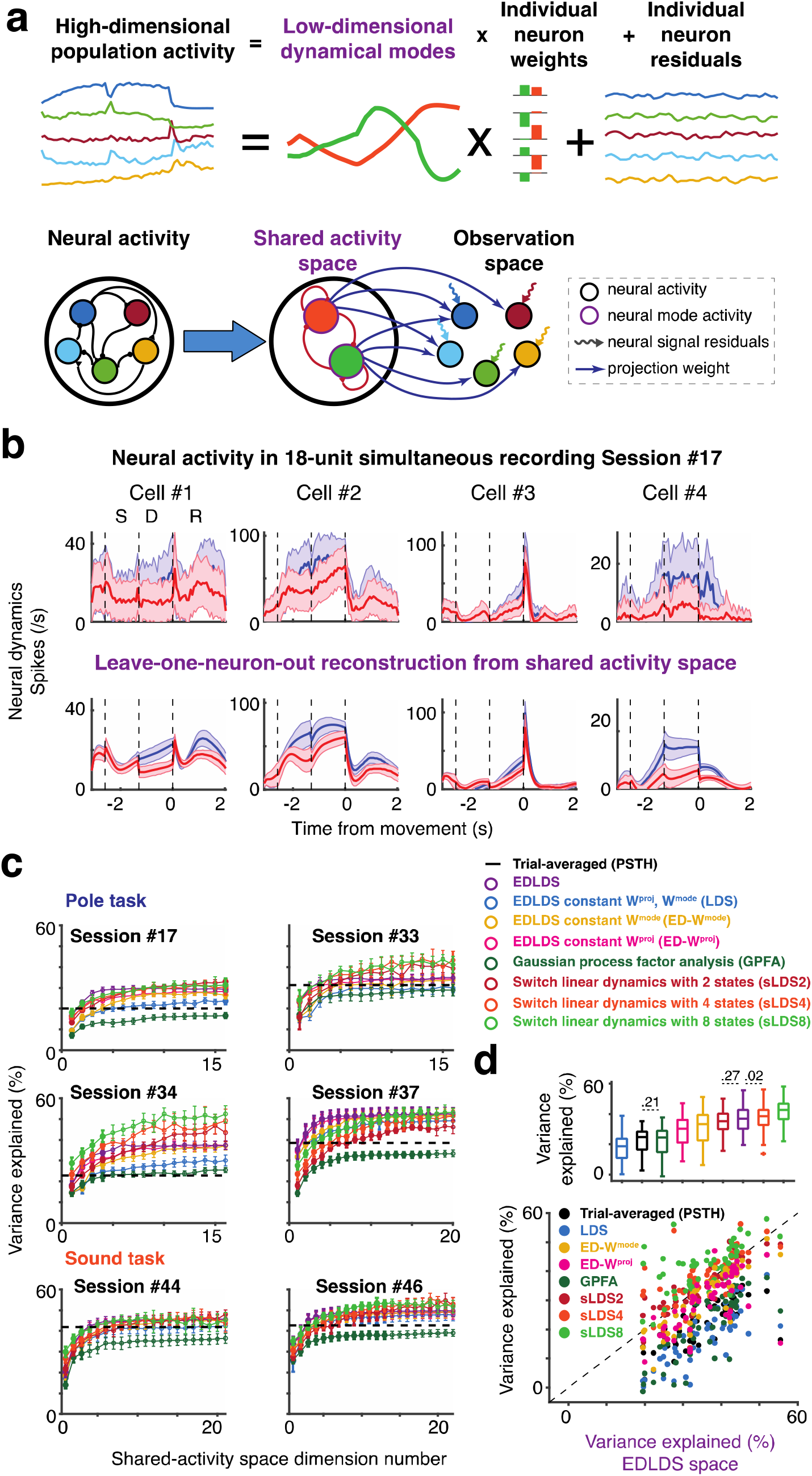
Latent dynamical system modeling of neural activity and the shared-activity space. **a.** Schematic description of time-varying linear dynamical systems model. Top: population activity is decomposed into a few activity modes shared across neurons. The activity of each neuron is described as a weighted sum of the shared modes and a single neuron residual activity. Bottom: alternative representation of model as a two-layer neural network. **b.** Model reproduces neural activity in a leave-one-neuron-out scenario; the activity of a given neuron is estimated from the dynamics of the other simultaneously recorded neurons. Top: Observed neuronal activity of four neurons randomly picked from Session #17, (ipsi trials red, contra trials blue). Bottom: prediction of neuronal activity on prediction data where the activity of that neuron is unobserved. Shaded area, STD across trials. **c.** Comparison of shared activity space fits for different models: epoch-dependent dynamical systems (EDLDS, purple, our model); three variations of our model - constant dynamical systems (LDS, blue), constant **W^mode^** (yellow) and constant **W^proj^** (pink); two simpler models - PSTH (mean activity in time for each trial type; black) and Gaussian process factor analysis (GPFA, dark green); and three variations of switch linear dynamical systems with 2 (red), 4 (orange) and 8 (light green) hidden states as a function of number of latent dimensionalities for 6 recording sessions (4 from pole tasks, 2 from sound tasks, note all data from simultaneous recordings, not pseudo-populations). **d.** Comparison of variance explained for different models for all recording sessions (n = 55). Scatter plot below shows each single session as a dot; boxplot above, shows across-session statistical summary; p < .001 for paired-comparisons between two models in boxplots without explicitly values.

Once we fit a model to a particular set of population activity, we measured the goodness of fit (R^2^) using a leave-one-neuron-out (LONO) approach^15^, predicting the value of the activity of a particular neuron when it is left out of the model fitting. If the dynamics of individual neurons are correlated and neural-mode dynamics have been well-estimated, it should be possible to predict the activity of any given neuron from the estimation of the neural mode dynamics derived from test trials in which the activity of that neuron was held out. Comparing the raw neural activity with its LONO estimation, we found that the dynamical model explained a large fraction of variability in the neural activity (*R*^2^ = .31, **Fig. 2b,** Session #17, N = 18 neurons, M = 4 modes). For most sessions, LONO R^2^ initially increased with latent dimensionality and then saturated (**Fig. 2c**). We compared variance explained among different configurations of our model, e.g., allowing only the latent dynamics to change from epoch to epoch, or allowing only the relation between latent dynamics and neural activity to change across epochs. We also compared our models to a large set of other models of varying complexity. For a reference value, we compared all models to a prediction based on the trial-averaged response (i.e., a neuron’s peri-stimulus time histogram, PSTH, for a particular trial type).

Incorporating the epoch-dependent matrices ***W***^*proj*^ and ***W***^*mode*^ yielded better fits compared to either of them being constant (**Fig. 2c**), consistent with the strong epoch-dependent change in dynamics. As expected, the EDLDS model outperformed simpler models (**Fig. 2d,** PSTH, Gaussian process factor analysis (GPFA), and LDS; p < .001) and performed only marginally worse than models that allowed switching at any time point (p = .27, sLDS2; p = .02, sLDS4; p < .001, sLDS8 and other models; sign-rank test of variance explained). In our hands the run time of sLDS models was 10-20 times longer, their fitting is more complex conceptually as well as algorithmically and there are hints of potential overfitting (i.e., switches at unexplained and inconsistent times; **Fig. S2cd**). Thus, we confirmed that the ESLDS model provided good fits to the population dynamics that are comparable to a true sLDS, despite its simplicity.

EDLDS models that provide good fits had a number of consistent properties. Nearly all sessions had a very long time constant during the delay epoch and in nearly all sessions the time constant during the presample epoch was the shortest of all epochs (**Fig. S2e**). Both of these are consistent with our expectation of a delayed-response decision-making task. In addition, the approximate dimensionality of the SAS was similar across sessions, (3 or 4, pole task; 6, sound task; **Fig. S2b)** despite the number of simultaneously recorded units spanned a fairly wide range.

### Decoding behavioral variability from the shared activity space

Having extracted single-trial dynamics in the SAS, we performed the most basic measure of their utility: decoding of behavioral variables. If the SAS judiciously pools neural activity across neurons and time, one expects better performance in decoding behavioral variability from the SAS than from the FNS. We first considered decoding trial type, training behavioral-choice decoders on either the FNS or the SAS at 300 ms before the onset of response. While both decoders successfully differentiated trial types (**Fig. 3a**), decoders running on the SAS were significantly more accurate (p < .001, sign-rank test, **Fig. 3b,** decoding based on different times within the delay period yielded qualitatively similar results).

**Figure 3.**
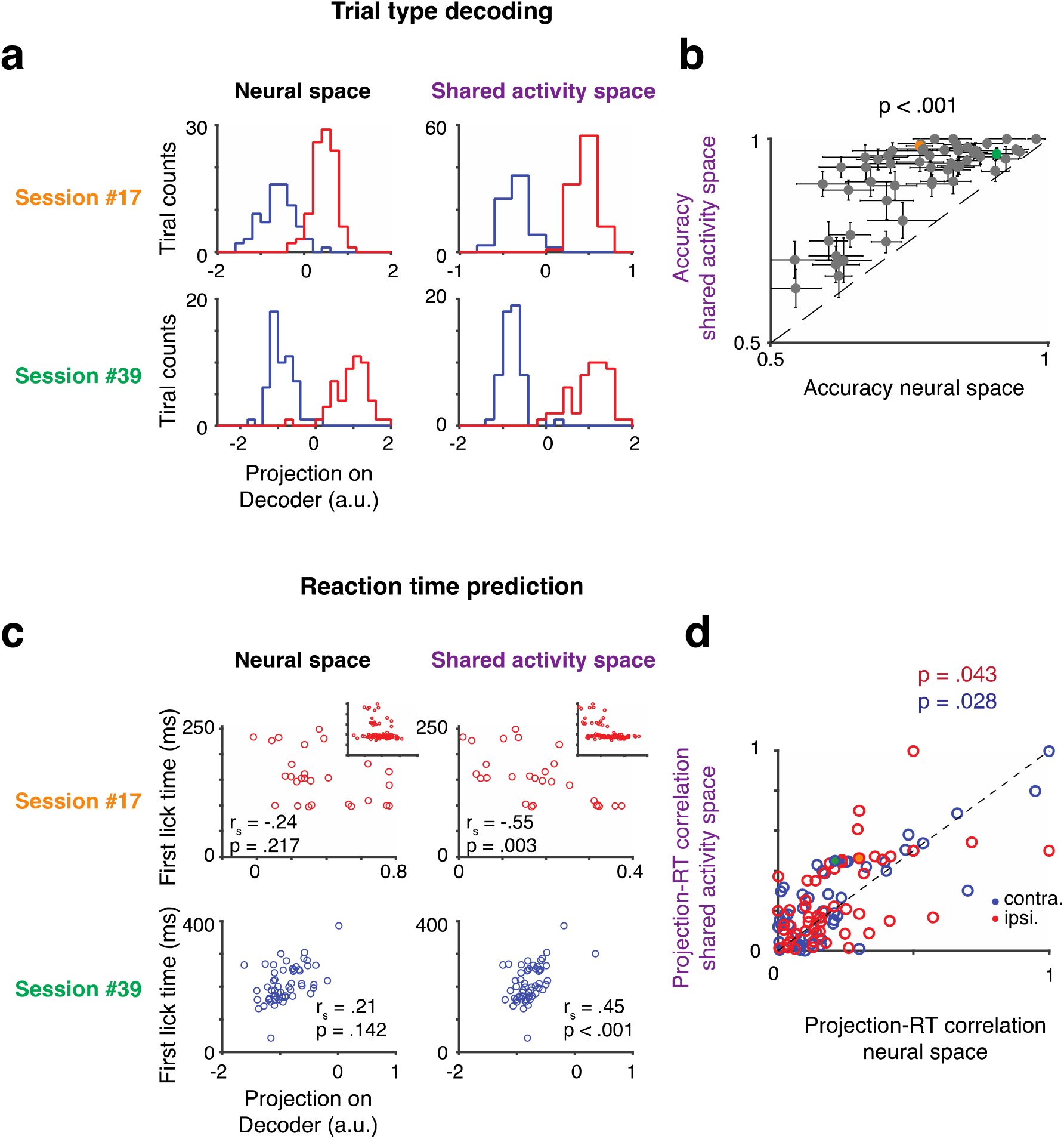
Neural dynamics in shared activity space correlates with trial-by-trial variability in behavioral reaction time and performance on single trials. **a.** Histograms of projection of all correct trials on decoder separately for each trial type (ipsi trials red, contra trials blue), for full neural space (left) and shared activity space (right). Session #17, upper; Session #39, lower. **b.** Accuracy of decoders based on full neural space (x-axis) and shared neural space (y-axis) across all sessions. Each circle represents a recording session. Error bar, SEM across trials. Session #17, orange circle; Session #39, green circle. **c.** Correlation between projection on decoder (x-axis) and reaction time (y-axis) for example session for decoder based on full neural space (left) and shared neural space (right). Top: Session #17, ipsi. trial types (blue), long reaction time trials (>100 ms), in main plot and all trials in inset. Bottom: Session #39, contra trial type (red), same convention. **d.** Scatter across all sessions of correlation between measured and decoded reaction time. Full neural space decoder correlation (x-axis) versus shared neural space decoder correlation (y-axis). For each session only one trial-type is considered. Each circle represents a session. Color of circle represents trial type; p-value presents significance that correlation is strong in SAS. Session #17, ipsi. trial type, orange circle; Session #39, contra. trial type, green circle.

Since trial-type prediction was already quite accurate in the FNS, we considered more subtle predictions, such as attempting to explain the variability in the timing of movement^7,33,34^. We performed this prediction based on the single-trial projection of population dynamics on behavioral-choice decoders. Projection values based on the SAS were significantly more correlated with movement onset than those based on the FNS (**Fig. 3c;** top inner, Session #17, ipsi. trials; FNS, left, r_s_ = -.31; p = .001; SAS, right, r_s_ = - .47; p < .001; bottom, Session #39, contra, trials; FNS, left, r_s_ = .21; p = .142; SAS, right, r_s_ = .45; p < .001; across sessions, **Fig. 3d**). We note that in multiple sessions, reaction times were a mix of short responses (100 ± 3 ms for the first lick) and longer (>200 ms, e.g., a long-tail distribution, **Fig. 3c,** top). In such sessions, considering short and long responses separately, long reaction times were predicted accurately in the SAS (**Fig. 3c,** top; Session #09; reaction time >100 ms; FNS, left, r_s_ = -.24; p = .217; SAS, right, r_s_ = - .55; p = .003) but short reaction time trials were not well predicted by either the FNS or SAS (p > 0.05). Comparing across latent space models, we found our ESLDS model outperformed most models in both trial type and reaction time decoding (**Fig. S3**). One potential caveat is that inferior performance in FNS could have been caused by poor model fitting, since the larger dimensionality of FNS forces decoders based on it to have more parameters. Therefore, we experimented extensively with regularization of FNS, yet the EDLDS-SAS performance was superior across all regularization parameters and methods^35^ (**Fig. S4**). Moreover, other models we compared had a similar dimensionality, thus did not require more regularization, yet still yielded inferior performance to the EDLDS-SAS. Notably, all the information to make predictions was in the FNS; the EDLDS and other SAS models are fit without additional side information. These models make specific assumptions regarding an underlying model for the dynamics, however, which allows them to perform effective denoising of signals and thus yield better predictors^7^ (see also Discussion).

### Long time scale states inferred from shared-activity space: continuous single-trial signals across epochs

Successful performance of the delayed task means that we should expect to find that trials from one trial type are consistently different from trials from another condition during the period from the sample period to the delay period. Beyond this minimal organization of population activity, it is not clear whether further levels of organization, such as a consistent relationship between trials from the same trial-type across time, can be discerned. Should such organization exist, it might reflect additional internal states that can be quantified, tracked, and potentially related to behavior^8,25^. This does not appear to be the case in FNS. Visualizing the projection of the first two principal components of the raw population activity across behavioral epochs, the trajectories of individual trials cross and mix (**Fig. 4a**). In contrast, the same type of projection performed on the SAS trials reveals an orderly relation within trial types (**Fig. 4b**). Namely, the position of a trial in the SAS extends throughout a single trial and systematically differs from trial to trial. Therefore, ALM population activity maintains a neural correlate of single trial identity beyond what is dictated by the task (since trials within the same trial-type are identical in terms of the task).

**Figure 4.**
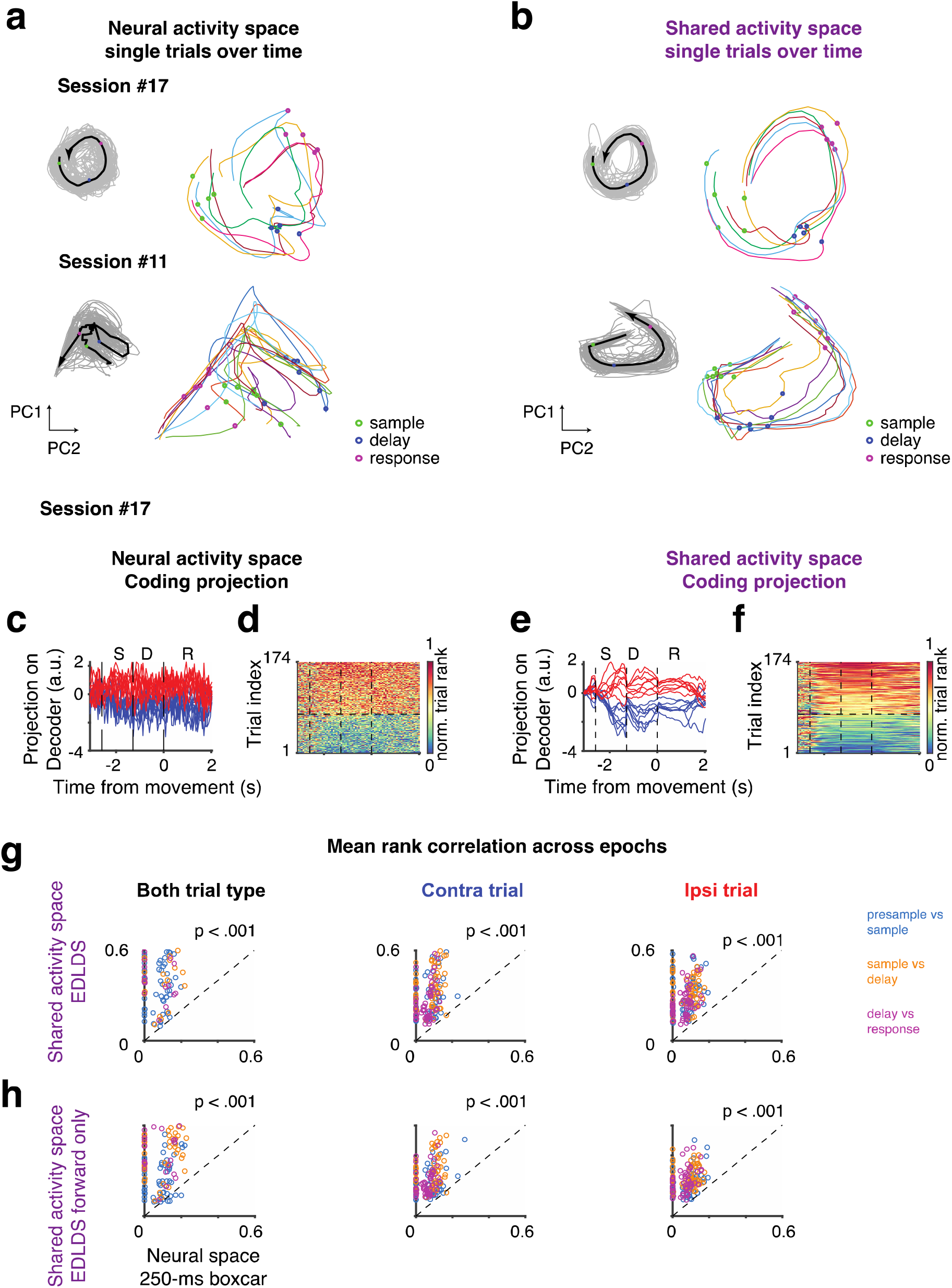
Maintenance of orderly internal representation of trial identity in the shared-activity space **a-b**. Neural dynamics spanned by first two principal components (contra trials, Session #17, top; contra trials, Session #11, bottom). Thin lines, single-trial dynamics; black thick lines, averaged dynamics across trials; circles, onset of behavioral epoch: sample (green); delay (blue); response (red). **a,** full neural space; **b**, shared activity space. Full neural space dynamics shows sharp transitions from one behavioral epoch to the other and random order on single trials. Shared activity space dynamics show smooth transitions in time and strongly ordered dynamics of single trials, where the inner trials (relative to mean) stay inner across time and behavioral epochs. **c-f.** Neural dynamics projected to instantaneous coding directions. **c.** Traces of single trial projections to the trial-types coding direction in full neural space. **d.** Rank of all trials across time. Data sorted according to average rank in sample-delay epochs. **e-f.** The same as **c-d,** except for those in shared activity space. **g.** Scatter of temporal correlations of trial identity across adjacent behavioral epochs in full neural space (x-axis) and shared activity space (y-axis). Each dot represents a session, color indicates transition type: between pre-sample and sample epochs (blue); between sample and delay epochs (orange); between delay and response epochs (purple). **h.** the same convention as **g.** except for forward-only pass EDLDS fit.

What is the nature of this correlate of single trial identity? We quantified it by tracking neural activity across time in each single trial and considering the relative relation between trials. Namely, we analyzed the projection on decoders of trial type since we and others have found these to be highly predictive of behavior^11^. Given the varying levels of neural activity across time points, we simplified the analysis by transforming the projection from its raw value to a relative rank across trials (either combining both trial-types or separately per trial-type). In FNS, the separation between trial-type was consistent across time, but there was little consistency of the rank within trial-type (**Fig. 4cd**). In contrast, for decoders built from the SAS we found that single trial projections both were separated across trial types and maintained single-trial ranks across time (**Fig. 4ef**). That is, a high-rank trial (high value of the projection on the decoder) tended to maintain a relatively high rank for the rest of the trial. Even when averaging activity across an entire epoch to improve signal-to-noise the single trial rank correlations across time were significantly higher in SAS than in FAS across all epochs (p < .001, **Fig. 4g**). This was not simply due to temporal smoothing in the SAS; decoders based on smoothed single trial activity had weaker single trial rank continuity (**Fig. S5c**). Nor was this effect solely due to de-noising by pooling across neurons; we found single trial rank consistency in the EDLDS-SAS was also superior to factor analysis, (**Fig. S5d**), implying that interaction among latent modes is related to the continuity of dynamics. Comparing across models, the EDLDS model shows a stronger consistency of single-trial rank than most other models (**Fig. S6**).

We were concerned that the increased consistency in rank might stem from the non-causal nature of fitting latent dynamical system models. This concern arises because these models infer the latent state at a given time point based on all time points both in the past and future. We addressed this concern by multiple controls. First, even when we ran the EDLDS model in a strictly causal mode (i.e., using past time points only to infer the current latent state, typically referred to as forward-pass) the rank was significantly more consistent across time than in the FNS (**Fig. 4h,** p < .001). Second, if rank continuity was merely a property the model imposed on the data, then it should be only weakly dependent on structure found in the data. In contrast, we found that shuffling neurons across trials within trial-type, i.e., taking the activity for each neuron and replacing it by the activity from the same neuron but from a different trial of the same trial type, significantly reduced rank continuity (p < .001, **Fig S7**). Thus, we conclude the persistence of the rank of a trial across epochs is an underlying property of the data that is revealed by the de-noising offered by the EDLDS model, and not an artifact of the model.

Next, we asked whether longer time-scale signals, on the order of tens of seconds or more, were discernible in the structure of population dynamics. We hypothesized that such signals might be revealed by similarity in the rank of activity in temporally adjacent trials (e.g., the next and previous trial). We found hints of such structure in the FNS by visualizing the rank of each trial across time and trials (**Fig. 5a**), and clear signs of rank similarity across multiple trials in the SAS (**Fig. 5b**). Once again this cannot be an artifact of the model since the latent state in the EDLDS is estimated separately for each trial and therefore the non-causal nature of the model is only within trial. To quantify these observed slow dynamics across trials, we computed the linear correlation between the temporal order of the trial (e.g., first trial, last trial) and the relative rank of its activity (within the same trial-type). Performing this analysis separately for each epoch, we found rank consistency within the SAS (*r* = .17 ± .02, contra, trial; *r* = .20 ± .02, ipsi. trial, mean ± sem, for hundreds of trials in every session) that was significantly stronger than in the FNS (**Fig. 5c,** p < .001 for all epochs) or other models (**Fig. S8a**). Interestingly, simpler models (i.e. LDS, GPFA, EDLDS) better revealed long-time scale dynamics than the full switching models (sLDS), perhaps because overfitting of sLDS model to short time-scale dynamics obscures longer time-scale correlations.

**Figure 5.**
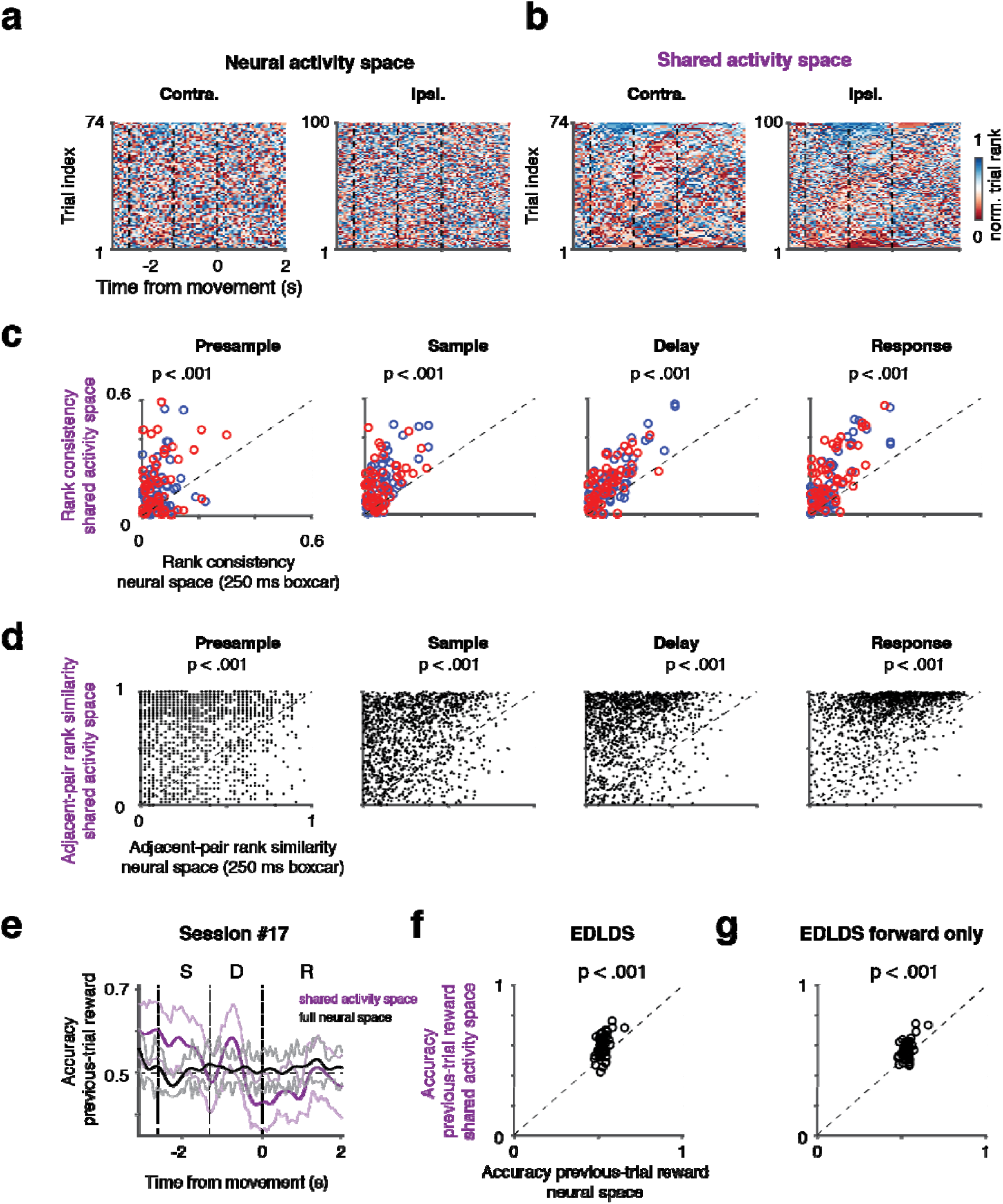
Long time-scale structure of population activity revealed in shared activity space. **a.** Heatmap of relative rank of population activity in full neural space for each trial type (74 contra, 100 ipsi trials). Rank at each time point was normalized to the maximum number of each trial type. **b.** The same plot as a. for activity in shared activity space. **c.** Comparison of rank consistency across trials based on the averaged activity within each behavioral epoch in shared activity space (y-axis) versus full neural space (x-axis). Blue, contra.; red, ipsi. trials; each circle represents a single session. **d.** Comparison of the rank-dynamics (change of within-epoch rank, see Methods) consistency between two adjacent trials (1344 pairs across 55 sessions) in shared activity space (y-axis) versus full neural space (x-axis). **e.** Decoding accuracy whether a previous trial was rewarded for decoders based on shared activity space in purple, for full neural space in black; shaded area, SEM across trials. **f.** Scatter of average decoding accuracy over pre-sample period for decoders based on full neural space (x-axis) or shared activity space (y-axis). Dash line, diagonal line. **g.** The same convention as in (**f)** except comparing to shared activity space based on forward-only EDLDS fit (y-axis).

We next considered an additional form of long time-scale structure: consistency in the way relative rank changes within an epoch (in contrast to the consistency in the average rank of a trial considered above). To clarify this type of consistency, relative rank within epoch would differ if in some trials activity ramps up during an epoch and in others it remains flat. This is because relative rank changes little when activity is flat, but it rises in later parts of the epoch when activity ramps up. We refer to the similarity of within-epoch rank changes as rank-dynamics consistency. We quantified rank-dynamics consistency by correlating the relative rank within an epoch between two trials:

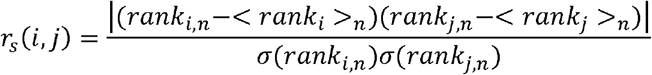

where *r_s_*(*i,j*) is the rank correlation of trial *i* and *j* in rank space for epoch *n* in the same trial type; <·>_*n*_ is an average over times in epoch *n*; and *σ*(·) is the standard deviation. We compared this quantity on neighboring pairs (*j* = *i* + 1) of the same trial-type only, which occur in a randomized trial label design, albeit infrequently (n = 1344 pair across 55 sessions). We found that rank-dynamics similarity was significantly stronger than the similarity found in the FNS (**Fig. 5d;** p < .001 for all four epochs). Comparing across all other models we found that the rank-dynamics similarity in the EDLDS was stronger than all models in nearly all behavioral epochs. Furthermore, all other models had at least one epoch that did not show significant rank-dynamics similarity (**Fig. S8b**).

We hypothesized that longer timescale features of the dynamics may relate to recent behavioral history. Indeed, we found that we could reliably predict the previous trial’s outcome^27^ (inter-trial intervals vary from 8 s to 20 s) during the presample epoch from the SAS, but not the FNS (**Fig. 5e-f;** other model fits, **Fig. S9a**). This was similarly true for other behavioral variables in previous trials, such as choice^36^, early lick, and stimulus^37^ (**Fig. S9**). The even longer timescale dynamics (tens of trials **Fig. 5b)** may relate to more difficult to track changes in behavioral state such as arousal or thirst.

In summary, we revealed rich structure in ALM dynamics using analysis based on the EDLDS latent space. Neural activity evolves in a manner such that an individual trial can be robustly tracked from sample epoch through delay epoch to response. Moreover, the dynamics of population activity change slowly over timescales of many seconds, with the dynamics seen in a particular trial more similar to the dynamics of a temporally-adjacent trial than a trial at some other time point, presumably due to the influence of a slow changing internal state. Finally, analysis in the SAS allowed explicit decoding of longer timescale behavior.

### Error trials

Given the unexpected level of structure in the neuronal population dynamics, we hypothesized that this structure might be relevant for understanding even complex behavioral patterns such as “error” trials where an animal makes an incorrect action. Analysis of error trials can lead to insight into the underlying computations^22^, for instance, they offer an opportunity to view a break in the typical, learned, association between stimulus and action. Yet analyzing such error trials is challenging because they are rare and the diverse causes for errors likely drive variable dynamics. Here, we compared the structure of dynamics in correct and error trials in the SAS. We hypothesized that the structure of dynamics differ grossly between correct and error trials^38^, such that error trials would be marked by a failure of the latent dynamical system model derived from correct trial data. Surprisingly, we found that a shared activity space built from correct trial data alone successfully predicted substantial variability in a LONO test during trials (**Fig. 6a;** captured 52 ± 7% variance of that using mean error-trial activity in two trial types, mean ± std.). This indicates that similar neural dynamics underlie the formation of correct and error trials, as in drift-diffusion models^39^. Additionally, although the fraction of explained variance was smaller in error trials (p < .001; **Fig. 6b**), its value correlated strongly with the explained variance in control trials. This indicates that the better the fit for the regular dynamics, the better the fit for the error-trial dynamics (r_s_ = .91, p < .001), again in line with the notion that qualitatively similar dynamics underlie correct and error trials (**Fig. S10a,** fit for other models).

**Figure 6.**
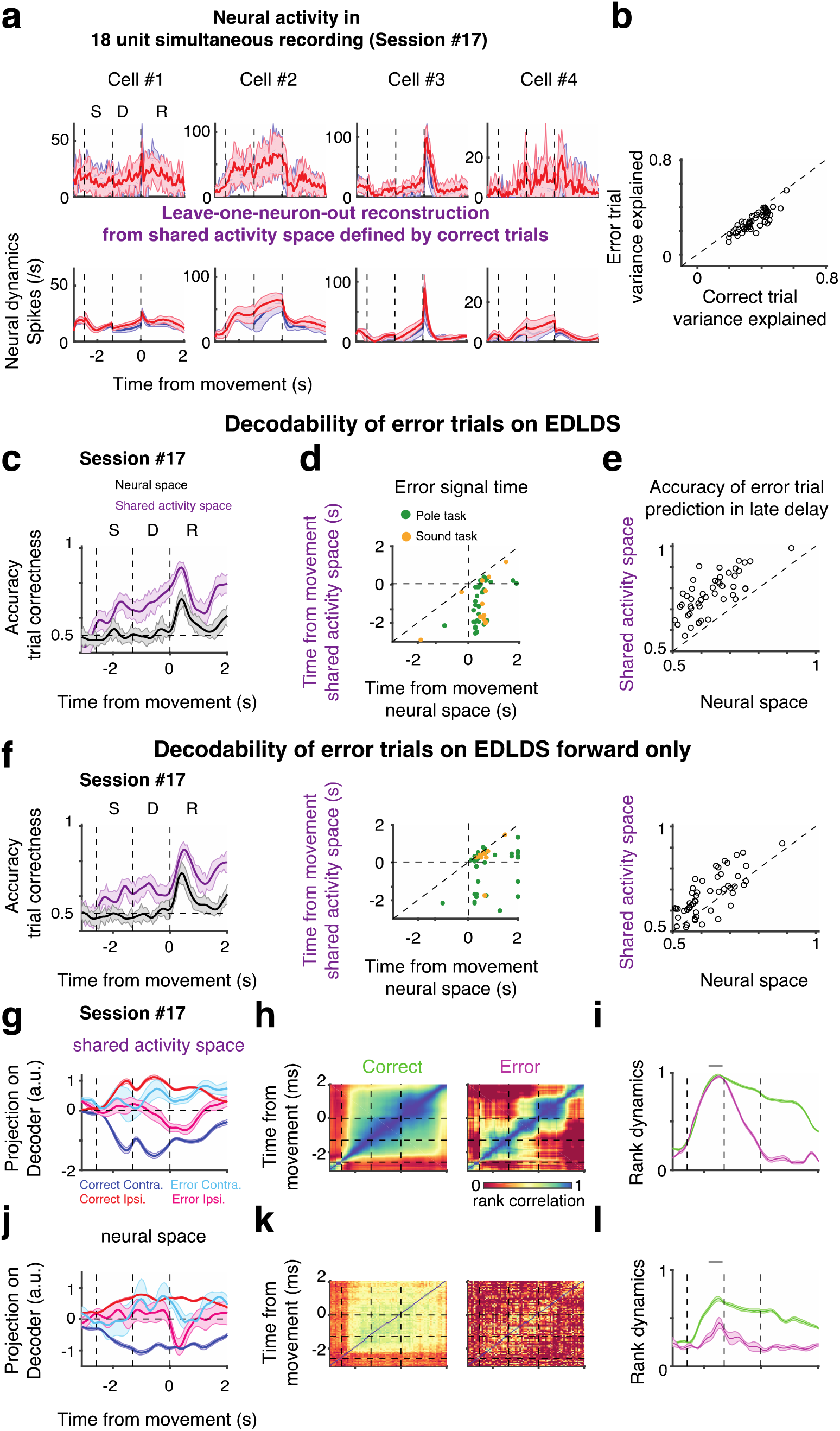
Error trials exhibit similar population dynamics to correct trials in the shared activity space. **a.** Neuronal activity in error trials can be explained by a dynamical system model latent space estimated from correct trial data only. Top: observed neural activity in error trials (cells identical to those in Fig. 2b); bottom: prediction of the neural activity using leave-one-neuron-out methods based on a shared activity space fit using correct trials. Shaded area, STD across trials. **b.** Fraction of variance explained by shared activity space model for error trials in different sessions (y-axis) scattered against fraction of variance explained by model on correct trials. **c.** Decoding accuracy of trial-correctness for decoders based on shared activity space in purple, for full neural space in black; shaded area, SEM across trials. **d.** Scatter of time in which decoding accuracy reaches a selected threshold (.65) for decoders based on full neural space (x-axis) or shared activity space (y-axis). Green, pole task, delay starts at −1.3 sec; Orange, sound task, delay starts at −2.0 sec. **e.** Scatter of decoding accuracy of trial correctness at late delay. Across sessions, decodability of trial correctness at late delay is high in SAS. Each circle corresponds to a session in **b, d** and **e. f.** the same convention as those in **c.-e.** except shared activity space is computed based on EDLDS forward-only pass (causal model). **g.** Plot of neural dynamics in shared activity space projected on decoders of trial type for correct (blue, contra.; red, ipsi. trials) and error trials (cyan, contra.; magenta, ipsi. trials). Bold lines, averaged activity; shaded areas, sem. **h.** Heatmaps of rank similarity in shared activity space across time averaged over correct (left) and error (right) trials. **i.** Similarity of instantaneous rank to that in the late sample (−300 ms to zero ms at the onset of delay, gray bar) in shared activity space averaged over correct (green) and error (magenta) trials. **j-l.** the same convention as those in **g-i.** respectively, but for neural dynamics and its rank in full neural space.

If the structure of population dynamics is similar across both correct and error trials, what makes a trial an error trial? Knowing what is the behavioral contingency, one can spot error trials in motor or decision areas by neurons having a response more similar to the opposite behavioral contingency^22^. However, this schematic description of flipping of responses does not always capture the dynamics observed in many neurons and brain areas^38^. Accordingly, there is deep interest whether error trials can be revealed directly by a more challenging decoding problem: decoding whether a trial was correct or not without access to its behavioral contingency (lick left or right). This decoding was far more effective in the SAS than in the FNS (**Fig 6c-e;** p < .001). In the SAS, accurate decoding of error started early in the delay period (**Fig. 6c;** Session #17), whereas in the FNS there was no successful decoding of error until the delivery of reward (**Fig. 6e**). Across sessions, decoders were able to decode errors significantly earlier in the SAS (p < .001; **Fig. 6d**). Furthermore, these results hold true when using the forward-only fit (p < .001; **Fig. 6f**), implying the results were not artifacts of the non-causal nature of the EDLDS model. Finally, the EDLDS model outperformed all other models in early predictions of error trials (**Fig. S10bc**). Taken together, these results underscore the utility of our approach.

Comparing across sessions that had sufficient error trials for decoding, we observed that the temporal profile of error decodability varied substantially from session to session (**Fig. S11**). We explored this issue using two approaches. First, we compared the projection on decoders of trial type for correct and error trials, then we inspected the rank-consistency we observed before. The average dynamics of error trials had diverse patterns, sometimes showing intermediate values between the trial conditions and sometimes showing a flip in the response to the other trial type (**Fig. S11b**). The timing of the divergence in error trial dynamics to the direction opposite to correct trials was typically consistent with the previous decoder results (**Fig. 6g**). When analyzing rank consistency, we lacked sufficient numbers of error trials to use within-trial rank as in previous analyses. Therefore, we used a rank measure that incorporates both trial types (mixing between and within group rank). We found clear disruption in across-epoch rank-consistency in error trials (**Fig. 6h**). Specifically, we compared the rank of trials at each time point with a reference rank calculated around the transition between sample and delay (−300 ms to 0 ms relative to delay onset, qualitatively similar results were found for other time windows around this period). We found that rank dynamics starts to diverge between error and correct trials after the end of the sample epoch, consistent with the timing inferred from the other analyses (**Fig. 6i**). We cannot rule out that the different temporal profiles of error decodability is not the product of differential sampling of neural selectivity across sessions, however, which cannot be directly controlled. Although we did not find any consistent patterns, there were sampling differences between sessions. For example, Session #17 has 7 trial-type selective neurons (among 18 units) from sample to delay epochs, and error trial decodability held continuously across sample-delay epochs. In contrast, Session #13 (**Fig. S10)** had one trial-type selective neurons in the sample epoch and 4 other neurons (among 11 units) in the late delay, and error trial decodability broke down in the middle of delay. Performing the same analyses in the FNS (**Fig. 6j-l**), we found the divergence of error trials still agreed with the FNS decoder results, but the rank-consistency was too noisy to reliably compare to the timing of significant decoding (results for other sessions, **Fig. S11**). In summary, the de-noising properties of the SAS allowed more fine-grained analysis of error related signals, with the inferred timing of hypothesized error generation consistent among multiple measures.

## Discussion

Population dynamics in neural circuits are rich and highly heterogeneous, with large variability in selectivity and response patterns both through time within a trial and across trials. Thus, accurately describing the underlying computations and their relation to behavior is challenging. Here, we revealed stable, ordered internal representations that bridged these heterogeneities by applying latent dynamical system models to the epoch-dependent nature of the observed dynamics. These models can serve as proxies for internal states and more accurately explained behavior in both correct and error trials.

### Latent space dynamical system models of neural population activity

Our EDLDS model accurately predicted multiple features of behavior such as an animal’s choice on single trials and its reaction times. Although the shared activity space has less information than the full neural space, it delivered superior performance from the full neural space in predicting multiple behavioral features. Two considerations may explain the superior performance of the shared activity space. First, if the shared activity space captures most of the relevant information from the full neural space with fewer parameters, decoders trained on the shared activity space will generalize better than ones trained from the full neural space. Importantly, latent dynamical system models, such as EDLDS, capture information not only across neurons but also across and time, unlike static decoders. Attempting to capture all this information with no compression results in many hundreds, if not thousands, of variables: a 4-second trial subdivided into 50-millisecond bins yields 80 temporal bins for the activity of each neuron. Recording 20 neurons thus requires 1600 variables to capture the full information. These parameters must be fit based on the data from a few hundred trials which can lead to poor model fitting. In principle, regularization can alleviate some of these difficulties, but when a dataset’s correlation and noise structure are unknown, choosing effective, generally-applicable regularization methods is difficult^40^. In contrast, the EDLDS latent variables are lower dimensional and their values at a given time point naturally incorporates past information even when decoders have access to data from only a single time bin. This greatly reduces the number of parameters and improves generalization. The second consideration that explains SAS performance is the utility of imposing the assumption that population activity follows reliable dynamics which (if true) can alleviate the effect of measurement noise. Consider a system for which the dynamics must *apriori* obey some rules (e.g., gravity). If our expectedly noisy observations produce a trajectory inconsistent with the known dynamics (e.g., a ball observed to fly up), observational error must have occurred. Accordingly, one must assume that the true underlying state was not that observed but rather some other trajectory consistent with the known dynamics. Such a process is known as “de-noising”. For neural data, the rules of populations dynamics are not known in advance and must be inferred from data. Yet, given that spiking activity induces substantial measurement noise, especially for low firing-rate neurons, such as those in ALM, analyses that incorporate de-noising are likely to be more robust. Since the dynamics need to be learned from data, simpler (and therefore more robust) models of the dynamics are preferred, such as linear dynamical systems. However, if the model of dynamics is too impoverished, imposing these dynamics on the data will introduce bias instead of denoising. We hypothesized that adding event-driven switches to linear dynamical systems, generating the EDLDS model, will generate a rich, but still robust model of dynamics and thus allow effective de-noising even when the dynamics need to be estimated from the data. Our results demonstrate the EDLDS model’s ability to assimilate and de-noise data, making it especially apt for performing single-trial population analysis^14-18,24-27,31^, a major goal of systems neuroscience.

Our results also shed light on how diverse models of population activity capture neural activity and relate them to behavior, an important question given that most studies will likely employ only a single model to interpret neural dynamics. There are multiple desiderata of a model of population activity. A key measure of any model is its ability to fit the data, which can be readily quantified by predictions on held-out data. As expected, we find that more complex models perform better in explaining variance of held-out neural data, but these models didn’t necessarily explain behavioral variance better and revealed less underlying structure in terms of rank consistency (**Fig. 7**). Specifically, we observed many cases in which the inferred switches in the latent dynamics of an sLDS model were not easily explained (**Fig. S2cd**), perhaps because these are overfit irrelevant switches which would in turn cause disruptions in the rank-consistency of the latent states and obscure the structure of population activity. We also observed a stark contrast between how easily these models are to fit. Fitting LDS or our EDLDS models is conceptually and algorithmically simpler than sLDS models, and it is one or two orders of magnitude faster (in our hands, changing the computation time for a typical set of sessions recorded in a single animal from weeks in sLDS to a few hours in EDLDS). More generally, the difference between EDLDS and sLDS can be thought of as using supervised switches versus unsupervised inference of switches. In situations when a given behavioral event, in our case a change in behavioral epoch, is likely to lead to a switch in the dynamics, one should adopt a methodology that explicitly takes this into account as in the EDLDS. On the other hand, by leaving the precise switch time flexible, one can use the inferred time of switch as an informative measure regarding the single-trial dynamics, as in sLDS. Future work can develop models that combine both cued and discovered switching or approaches that allow fine-grained discovery of the precise timing of an expected switch. The dataset we use in the paper will become freely available and has several interesting and non-trivial properties (**Fig. 7)** that could make it a useful benchmark or reference point for testing new models.

**Figure 7.**
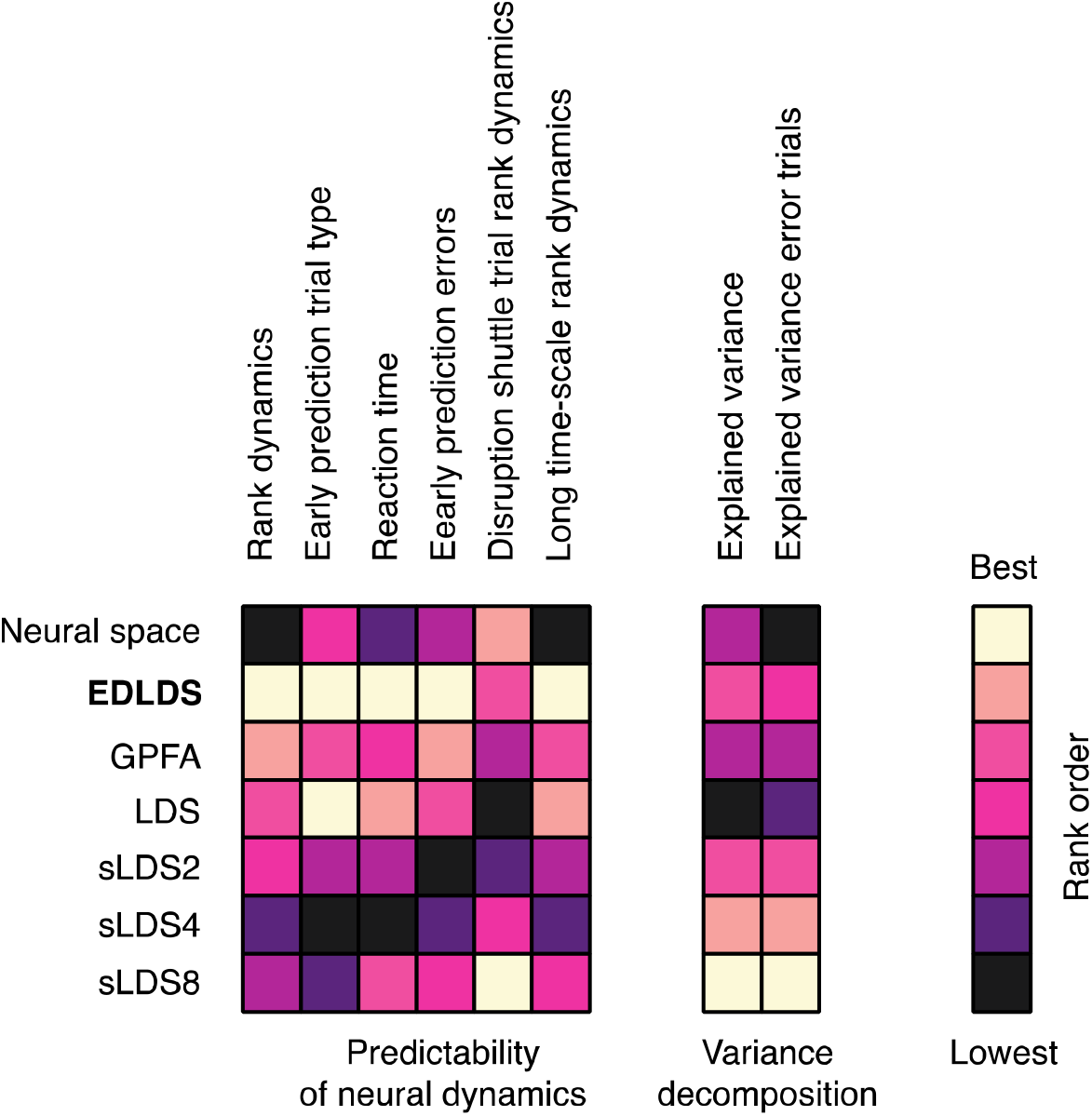
Summary of comparison across models. Results from Fig. 2 to Fig. 6 were compared across models and summarized into a pairwise comparison table. The comparison is divided into two categories according to the nature of the analyses. The first category emphasizes the predictability of behavioral and neural variability from different models. The analyses in this category includes: (1) rank dynamics: **Fig 4g, S5;** (2) early prediction of trial type: **Fig. 3b, S3a;** (3) reaction time: **Fig. 3d, S3b;** (4) early prediction of errors: **Fig 6d, S9b;** (5) disruption shuffle trial rank dynamics: **Fig S7;** (6) long time-scale rank dynamics: **Fig. 5cd, S8.** The second category emphasizes the variance explained by each model. This includes the explained variance for correct trials (**Fig. 2d)** and error trials (**Fig. 6b, S10a**).

### Error trial analysis

Error trial analysis can illuminate the nature of a neural representation^22,23,34^, but analyzing error trials is challenging due to their scarcity and the potentially diverse origins of errors (e.g., a mistake in sensory processing, a lack of alertness, low motivation etc.). Accordingly, error trial analysis stands to greatly benefit from latent space approaches that allow more powerful single trial analysis. However, the small number of error trials renders fitting a dynamical system model of them unfeasible. It has been argued that errors are caused by a strong change in the dimensionality of a representation^38^, and thus adopting the same underlying dynamic model from correct trials would not be appropriate. In addition, it is intuitively reasonable to expect that at least some forms of error, e.g. lack of alertness, would fundamentally affect the underlying computations and thus also the underlying dynamics, again rendering them distinct from correct trials. To our surprise, we found that the dynamical structure of the population activity is surprisingly similar in correct and error trials. This is important from a conceptual point of view as it provides evidence against models that assume dynamics are different in error trials, and for other models in which error trial dynamics are more similar to correct trial dynamics such as drift-diffusion models^39^ and some attractor models^41^. Practically this is important since most latent dynamical system models would not be reliably constrained by using the small number of error trials but can be fit with the number of correct trials.

Using the models fit on correct trials we were able to perform a quantitative comparison of neural activity in correct and error trials. Decoders based on the superior denoising by the EDLDS models robustly predicted neural signals associated with behavioral errors seconds before movement onset. Furthermore, we leveraged the additional structure in the organization of single trial dynamics revealed by our analysis, i.e., the consistency of single-trial rank across time within a trial, to infer both the existence of an error trial and the likely timing of its occurrence. Specifically, the latent representation of a single correct trial maintains its relative rank in time, but the relative rank of an error trial is disrupted at some point in error trials. Importantly, we stress that comparison of latent dynamics in correct and error trials goes beyond (and is complementary to) single neuron error-trial analysis, and is only possible at the population level^42^. Future experiments which allow independent analysis of whether and when an animal is likely to make a mistake, e.g., inferred inattention from pupil size, will be important to assess the advantages of finer-grained error trial information revealed by latent space analysis.

### Structure in neural dynamics and its relation to internal states

In trial-based designs, one can compare how behavior differs across diverse trial conditions to the cognate trial-averaged population activity. Similarly, one can perform decoding analysis to regress neural activity against measured behavioral variables such as the instantaneous position, or velocity, of an animal or one of its limbs. Such decoding analyses are robust for aspects of behavior that we directly impose, such as trial condition, or can reliably measure, such as position. Yet the animal has additional internal states that affect behavior such as alertness, motivation, hunger, and others that can only be partially deduced from observation, e.g., of pupil diameter. Moreover, it is likely that there are many additional states that cannot be dictated by experimental paradigms or measured by tracking the animal.

Yet these states don’t have to remain a black-box; assuming these states influence neural activity, it may be possible to identify or infer them from population recordings, e.g., from specific brain regions whose raw activity directly reflects the state or in subtle activity changes in other areas. Central decisionmaking hub areas (e.g. ALM, or other parts of frontal cortex; **Fig. S1g-i, S1m-o**) are key candidates for such areas where long time-scale dynamics might be revealed since they likely integrate many types of information and their activity is directly related to behavior^11^.

Long time-scale organization of activity is of particular interest both due to its temporal match with internal states that are thought to be long-lasting, and from the technical perspective of inferring structure in limited signal-to-noise sampling since sustained structure in single trials is unlikely to occur by chance. In principle, such structure might be observed from raw single neuron or population activity (particularly when behavior itself is correlated strongly with previous trial behavior^36,37,43^), but it may be obscured by the estimation noise induced by the spiking of neurons, especially in neurons with low firing rates. Here, we found strong evidence for such structure both across the seconds-long time scales expected from the nature of the task, i.e., the timescale of transfer of information from sample to delay to response period, and across longer, multi-trial, many-second timescales. This organization can be faintly observed in the raw neural activity, but the denoising offered by the latent state models strongly and significantly unmasks this structure. This reveals an additional level of organization in neural activity and a potential leverage point for understanding long timescale internal state dynamics in different brain areas and understanding longer timescale phenomena such as learning or motivation (thirst level, alertness etc.). Finally, in the future, advanced non-linear decoders might leverage this structure to perform superior denoising.

More generally, an elegant framework for the goal of neural computation is to move from a highly angled representation at the input level, such as pixel-level representations that represent a face at various angles and under different lighting conditions, to a more linearly separable representation in higher cortical areas^44,45^. Similarly, a key goal of single-trial population analysis is to reveal internal representations that maintain single trial information despite potential entanglement across time and neural space. Our analysis revealed that sharp changes in neural dynamics occur in a coordinated fashion across trials, such that a candidate neural signature of a particular trial’s identity is maintained across time. Although some orderly relations among trials have been shown previously, e.g., in large shifts of neuronal gain across trials^46^, here we utilize a dynamical latent space model to show that cortical activity exhibits an orderly, multi-timescale organization of its population activity.

The EDLDS model we introduced is a compromise between overly impoverished, fully linear models^30,31^ and more expressive models that are more difficult to fit^18^. Its power lies in its ability to discern features of population activity that are not obvious in the original recordings, i.e., the full neural space. And, while such features may not be discernible in all brain areas or all latent space analyses^8,25^, such approaches will become more and more important as we strive to study more demanding tasks.

## Authorship

Z.W and S.D conceived the project. H.I and N.L performed electrophysiological recordings. Z.W constructed the latent-space model. Z.W and S.D analyzed the data. Z.W, K.S and S.D wrote the paper with comments from the other authors.

## Acknowledgements

We thank Misha Ahrens and Xiao-Jing Wang for comments on the manuscript, Scott Linderman for the generous help with the switching linear dynamical system (sLDS) code (http://github.com/mattii/pyslds) and writing with us the code of sLDS leave-one-neuron-out estimation^31^. This work was funded by the Howard Hughes Medical Institute. K.S, S.D and N.L are funded by the Simons foundation collaboration on the global brain SCGB 325453; Z.W is funded by SCGB 542943SPI.

## Methods

### Behavior

Mice were trained to perform a delayed version of a two-alternative forced-choice discrimination task^11^. Mice reported the position of a pole (anterior or posterior) or the frequency of a sound (low or high) by directional licking (lick-left, ipsi trial; or lick-right, contra trial) after a delay period (sample period: 1.3 seconds, delay period: 1.3 seconds, pole task; sample period: 1.15 seconds, delay period: 2.0 seconds, sound task). Inter-trial intervals were variable, ranging from 8-20 seconds from the previous trial’s go-cue. In addition, the next trial was not initiated until at least 4 seconds had passed after mice stopped licking. Mice were water restricted. Delivery of water began immediately following the detection of a correct lick. Reinforcement was not delayed and water was consumed during subsequent licking. Mice were trained to a criterion of at least 70% correct.

### Electrophysiological recordings

Electrophysiological recordings were performed on the left-hemisphere anterior lateral motor cortex (ALM) using 32-channel NeuroNexus silicon probes (n = 19 mice) or 64-channel Janelia silicon probes (n = 20 mice). The details of electrophysiology and spike sorting were described in^10,12,13^. Regardless a few sessions were reported previously^10,12,13^, all sessions will be recompiled and released to crcns.org with codes (https://github.com/zqwei/TLDS_ALM_Data) that can recreate all figures in the main text.

We excluded trials with early licking. Cells were previously classified as fast-spiking interneurons and pyramidal cells in^10,13^. Recording sessions were chosen based on two criteria: each session must have more than 5 units and the number of correct trials should be more than double the number of units (8 sessions in 32-channel recordings; 47 sessions in 64-channel ones). Summary is shown in **Supplementary Table 1.** Session #17 was used as the example session throughout the main text. This session contained 18 simultaneously recorded neurons at depths 150 – 1000 um and was one of the two sessions composed entirely of pyramidal cells.

### Single neuron analysis

For all analyses, we binned neural activity using 67-ms discrete time window expect for when we explicitly compared to longer windows as a control in **Figs. 4g, S4c,** where we used 250-ms bins in 10-ms steps. To compare the single trial variability in different neural spaces, we computed *std*. of neural dynamics within the same trial type; except for in **Fig. S1g-i; m-o**, where sem. are presented.

To measure the dynamics of selectivity, we performed two-sample t tests with neural activity in each 67 ms discrete bins (**Fig. 1b**). We defined a neuron as monophasic if it had consistent polarity of selectivity (p < .05) for >335 ms (5 continuous bins), a neuron as multiphasic if it had a switch of selectivity (p < .05) with the periods of selectivity being at least 335 ms. The rest of the neurons were considered as nonselective neurons. For monophasic-selective neurons, we classified them into contra.- and ipsi.- preferring cells, according to the trial type for which they had higher activity (p < .05, two-sample t-test). Instead of correcting for multiple comparisons due to independent tests at each time bin we consider a neuron significant only if it had a significant value at 5 consecutive bins.

### Epoch-dependent linear dynamical system models and neural mode dynamics

Our Epoch-Dependent Linear Dynamical System (EDLDS) model, is an extension of a linear latent space dynamical system model^30^. In such models, the full neural activity, (***r***(*t*) ∈ ***R***^*N*^), is modeled as a projection up from a low dimensional neural-mode space, (***x***(*t*) ∈ ***R***^*M*^, *M* < *N*) by a projection matrix, **W**^proj^. The low dimensional modes evolve according to linear dynamics, **W**^mode^, with Gaussian innovations (**Equations 1 and 2**). In our model the model parameters are fixed in time but are allowed to be different for each experimental epoch, *s*. This model is a compromise between models in which there is only one fixed latent model, which have dynamics that are incapable of capturing the rich dynamics we find in ALM and models that allow switching at any time point^18^, which suffer from difficulties in estimation due to the combinatorial number of possible hidden states of transitions over time. Under this model, the joint probability of the data and latent modes is:

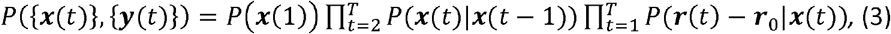

where 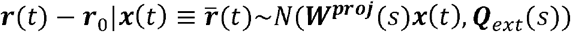, ***x***(*t*)|***x***(*t* – 1)~*N*(***W***^*mode*^(*s*)***x***(*t* – 1), ***Q***_*int*_(*s*)), and the initial state of the neural mode ***x***(1)~*N*(***x***_0_, ***Q***_0_).

The parameter set *Θ* included the amplitudes of external independent inputs 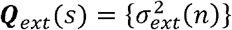 each neuron (1 ≤ *n* ≤ *N*; *N*, the number of neurons in simultaneous recordings), the amplitude of internal inputs 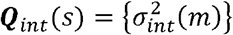 onto each neural mode (1 ≤ *m* ≤ *M*; *M* < *N*, the latent dimension), the latent connectivity, ***W***^*mode*^(*s*) ∈ ***R***^*M*×*M*^, and the projection, ***W***^*proj*^(*s*) ∈ ***R***^*N*×*M*^ for each epoch (i.e. pre-sample, sample, delay, and response), and those associated with the initial state of the neural mode ***x_0_***, ***Q_0_*** in trials.

The estimation of the EDLDS followed similar Expectation-Maximization steps as that in^14^. The expectation step was identical and the maximization step was:

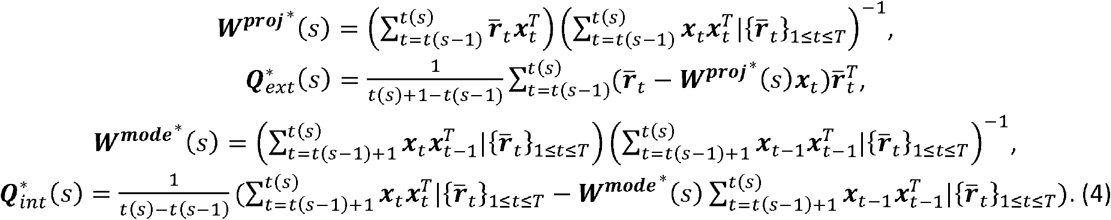

where *t*(*s*) presents the last time point of epoch *s* and *t*(0) = 1. Here we constrained the estimation of input matrices ***Q***_*ext*_(*s*) and ***Q***_*int*_(*s*) to be diagonal after each iteration of maximization. Further, we computed dynamics of the neural mode in the shared activity space as 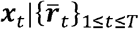 from the expectation step.

Model estimation was based only on correct trials (**Table S1**).

### Leave-one-neuron-out estimation and choosing dimensionality of the shared-input space

To determine the performance of model fits, we computed the variance explained using a leave-one-out procedure in cross-validation^15^. After fitting the model on training data, we take test data, remove the activity of the *i-th* neuron from the data, hence leave-one-neuron-out (LONO), and estimate the values of the shared activity space modes: 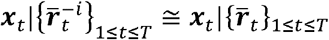 We then compute the expected activity of the *i*th neuron as 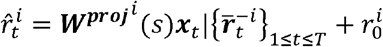, where ***W***^*proj*^*i*^^ is the ith row of the projection matrix ***W***^*proj*^. The explained variance,

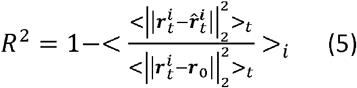

measured the goodness-of-fit (where || · ||_2_, is the *L*^2^ norm of the vector; <·>_*t*_, is an average over time and trials; <·>_*i*_ is an average over neurons).

To select the dimensionality of the shared activity space, M*, we performed LONO estimation of TLDS fit using 10-fold cross-validation for each possible dimension, 1 ≤ *M* ≤ *N* – 2 (**Fig. 2c**). The amount of the variance explained as a function of the dimensionality of the shared activity space first increased (corresponding to an under-fitting region) then saturated or even decreased (an over-fitting region). To avoid overfitting we picked the minimal dimension that reached a criterion of 90% of the maximum amount of variance explained. As a control we also compared results to the amount of variance explained in a model where either ***W***^*mode*^ or ***W***^*proj*^ was constant matrices (**Fig. 2c**).

As a comparison, we computed explained variance of a reference model, the mean-activity model, where we assume that neural activity follows a random fluctuation around its mean in each trial. Therefore, for each neuron, 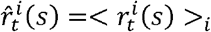, and index s stands for the trial type: correct contra, correct ipsi, error contra, or error ipsi trials. Computation of explained variance then follows **Equation 5.**

### Decoding analysis

To determine trial-type decoding we applied linear discriminant analysis (LDA) on neural dynamics grouped into 67-ms non-overlapped bins. The LDA decoder, ***l***_*t*_, was computed separately for each time bin, t, using correct response trials only. The neural dynamics projected onto the decoder was then computed as 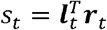, where ***r_t_*** is the vector of neural activity in time bin t. To avoid the ambiguity of sign in the decoder we assigned a sign to the decoder so that ***s_t_*** < 0 in contra and ***s_t_*** > 0 in ipsi trials. The same procedure was applied for decoders trained on the shared activity space. The correlation of projected activity across time, *t*, and, *t*′, was performed using Spearmen’s rank correlation, *r_s_*(*t, t*′) and that across epochs, *s*, and, *s*’, were as < *r_s_*(*t, t*′) >_*t*∈*s;t*′∈*s*_ (**Fig. 4g,h, S5**). Epoch coding decoders were based on the neural dynamics averaged within each behavioral epoch; trial-coding direction is based on the neural dynamics averaged from sample to response epochs.

To achieve robust estimation given the large number of neurons, we applied a sparse version of LDA ^35^, with normalization that ||***l***_*t*_||_2_ = 1 (where || · ||_2_, the *L*^2^ norm of the vector) for each time point t in a task.

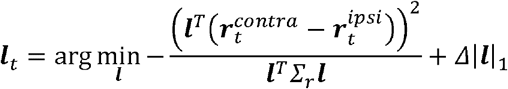

Two parameters were used to regularize sparsity of LDA coefficients, *γ* (*l*_2_-regularizer of covariance matrix of sample data 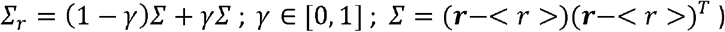 and *Δ* (*l*_1_ - regularizer of covariance matrix of sample data Δ|***l***|_1_). The optimization of these two parameters follows the standard Matlab procedure (Fig. S12) using cross validation (all trials were divided into training and testing sets), and the set of parameters minimizing the validation error were used in regularizing ***l***_*t*_ in final fit using all trials. In this case, the LDA coefficient should be close to zero if a neuron fired at a low rate or contributed little to coding trial type. We measured the similarity of the coding directions across time, *t*, and, *t*′, as 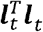. We did not observe any 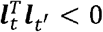.

### Decodability

The performance of trial-type decodability was computed based on the decoder projection in a 150-ms time bin at 300 ms before the onset of response using 10-fold cross-validation (**Fig. 3ab;** error bar, std.). We computed performance of trial-correctness decodability using two nonlinear decoders (**Fig. 4c; Fig. S7**), one based on a 2^nd^ order polynomial-kernel support vector machine (SVM; kernel function, 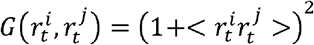; i, j, neural index; r, neural activity; <·>, average over trials), the other is based on quadratic discriminant analysis (QDA), based on instantaneous neural dynamics in discrete 67-ms time bins, using 10-fold cross-validation (shaded area, std.). The correctness signal onset time (**Fig. 4e**) was computed as the first time when performance of trial-correctness decodability was continuously > .65 for at least one behavioral epoch. We then compared the single-trial decodability in FNS versus that in SAS using paired t-test across trials.

### Reaction time correlations

We correlated decoder projection scores with reaction time to the first lick using Spearmen’s rank correlation. The decoder projection score was estimated by averaging over a 150-ms time bin at 300 ms before the onset of response (**Fig. 3cd; Fig. S3b**).

---

**Supplementary Figure 1.**
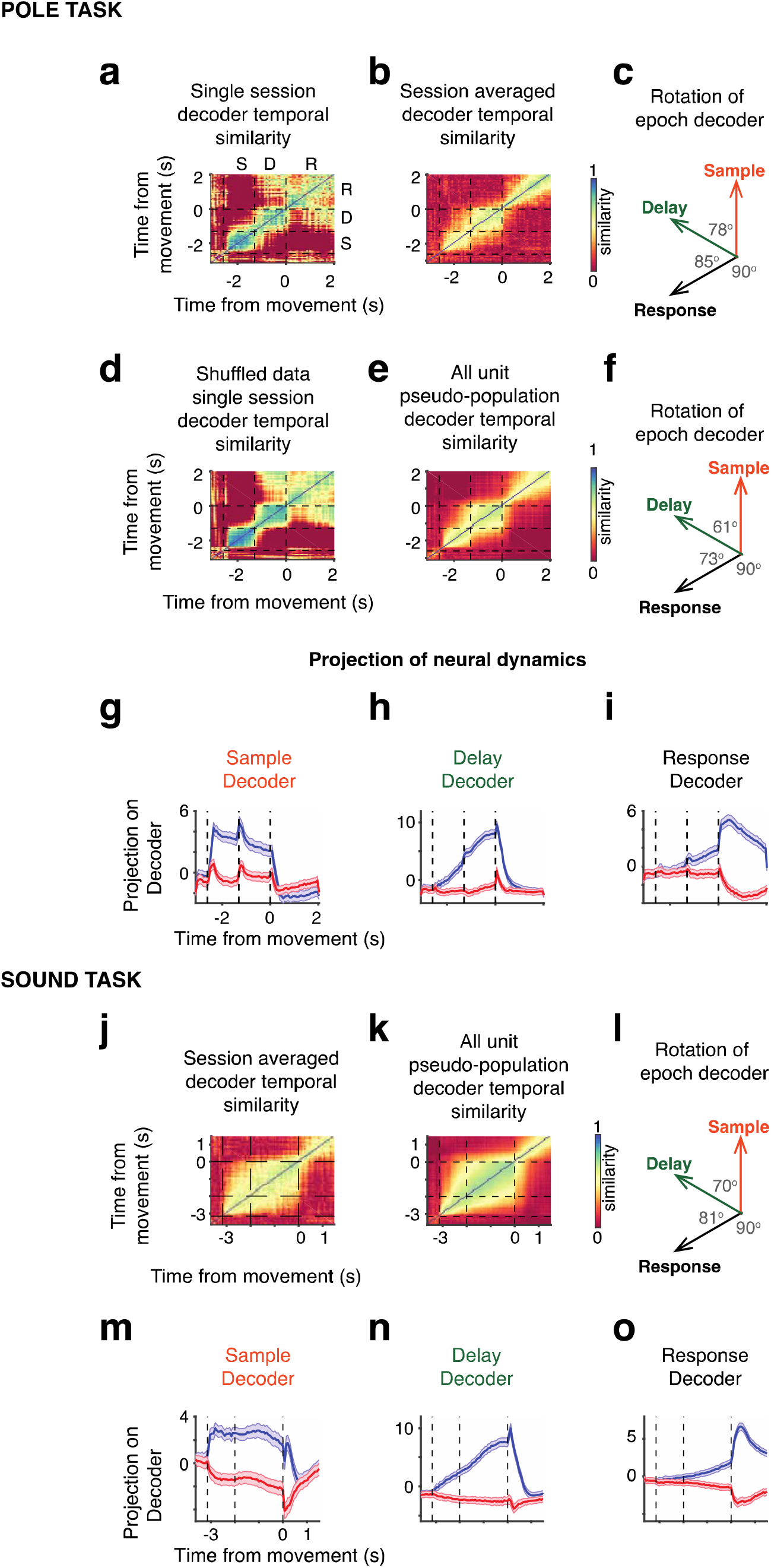
(Supporting figure for Figure 1) Rotation of coding directions and population dynamics in full neural space. **a-f.** Rotation of coding directions in full neural space. **a-c.** Similarity of instantaneous trial-type decoders across time. Color bar on right indicates correlation between decoders at different times. Coding direction is similar within epoch but different from sample-delay epoch to response, indicating the discontinuous dynamics of selectivity at population level. **a.** Single session analysis (n = 18 neurons; Session #17). **b.** Same analysis averaged across sessions (n = 40 sessions). **c.** Schematic of the rotation of decoders across epochs and value of angles between epoch-averaged decoders. **d.** Temporal correlation between instantaneous decoders across time similarity plot, same session as **a,** but with shuffled trials. **e.** All unit pseudo-population temporal correlation between instantaneous decoders (n = 1,743 units, randomly shuffled pseudo-combined trials). **f.** Schematic of the rotation of decoders across epochs and value of angles between epoch-averaged decoders for the all-unit pseudo population. **g-i.** Projection of the pseudo-population neural activity on trial type decoders based on activity averaged across each behavioral epoch: sample (**g**), delay (**h)** and response (**i**). **j-o.** The same convention as that in **a-i,** except for sound task.

**Supplementary Figure 2.**
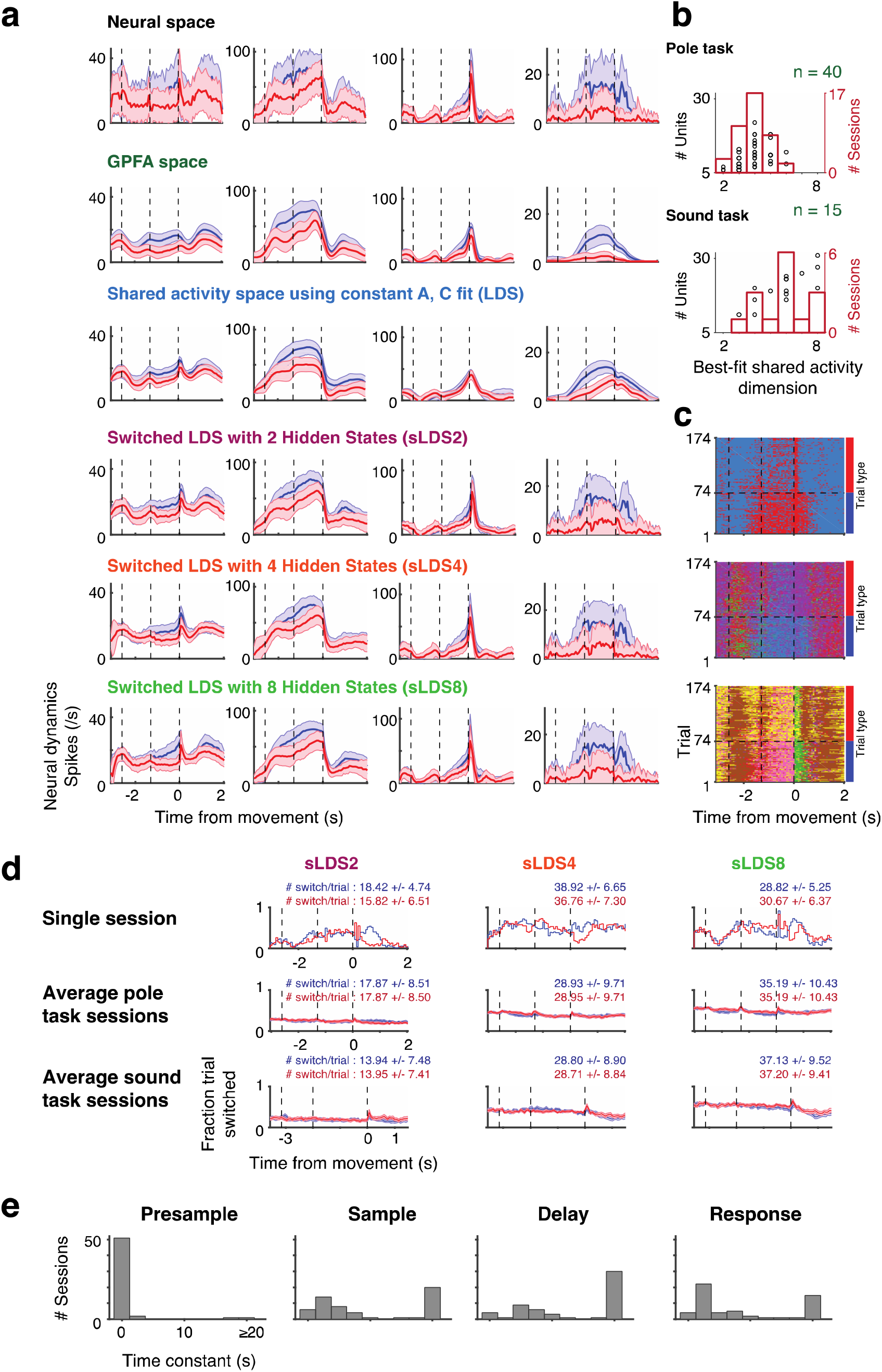
(Supporting figure for Figure 2) Neural activity fits on other models. **a.** Fit of the same neuronal activity in Fig. 2b using other models. **b.** Variance saturating dimension (VSD) of latent space model for all sessions using EDLDS fit. Plot shows each session as a black circle; x-value, VSD for that session, y-value, number of units. Red bars show number of sessions with a given VSD. Top, pole task; bottom, sound task. **c.** Fit of hidden states in different sLDS models. Top, sLDS2 (2 hidden states); middle, sLDS4 (4 hidden states); bottom, sLDS8 (8 hidden states). Each color present a unique state in heatmaps, and fit allows the switch at any time during a trial. The average number of switches in each trial type were reported in d. **d.** Fraction of switch happens uniformly in time. **e.** Distributions of time constants of fits in EDLDS model for different behavioral epochs (tailed at 20 s). The time constants is computed based largest of eigenvalue of ***W***^*mode*^, where 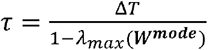; *λ*_*max*_(***W***^*mode*^) for each epoch in a session is reported in Table S1; Δ*T* = 67 ms is the time bin in our fits; for ***W***^*mode*^ > 1, we consider *τ* > 20 s.

**Supplementary Figure 3.**
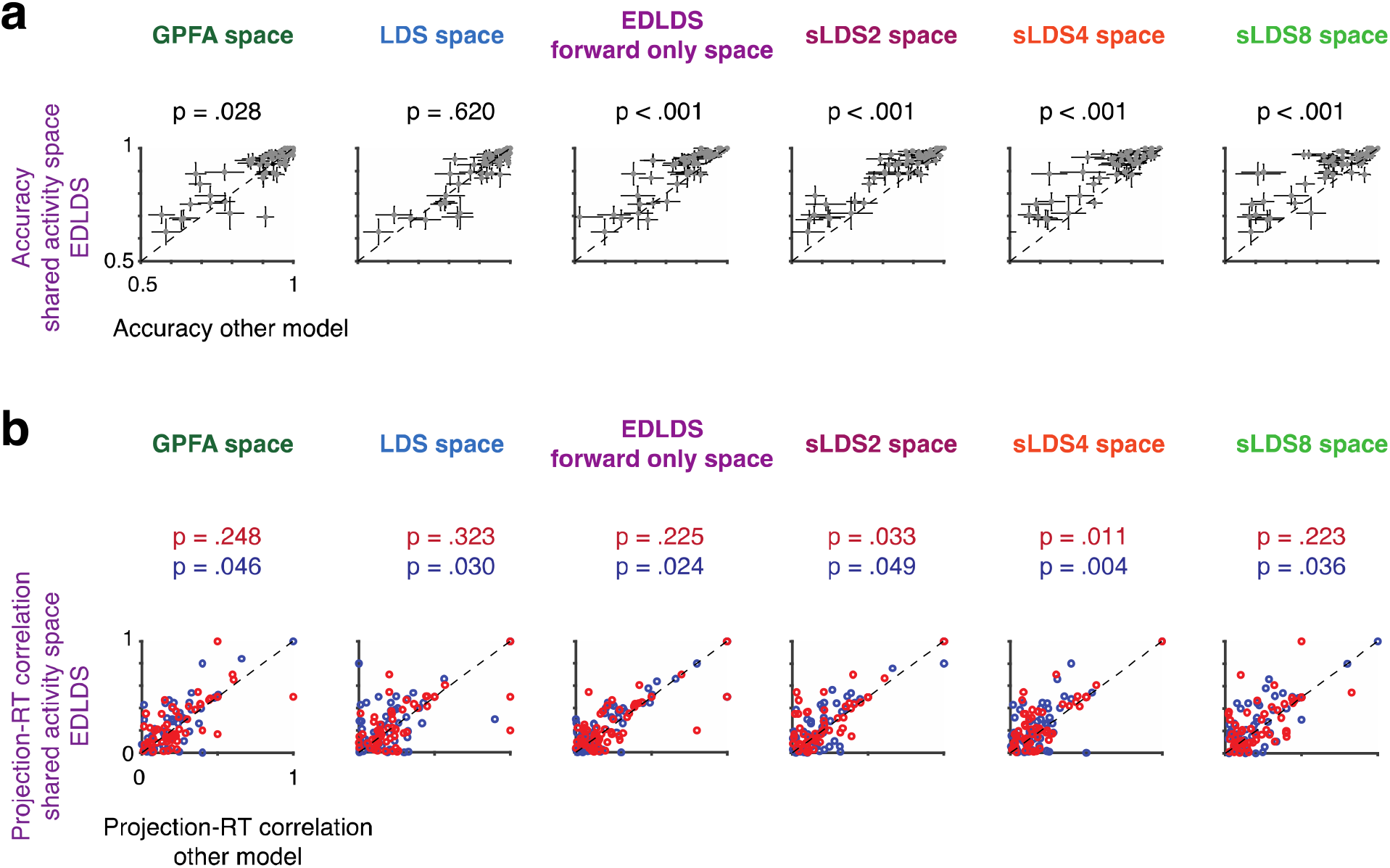
(Supporting figure for Figure 3) **a.** Following the same convention as that in Fig. 3b except for other models. **b.** Following the same convention as that in Fig. 3d except for other models. Both p-value is reported using sign-rank test.

**Supplementary Figure 4.**
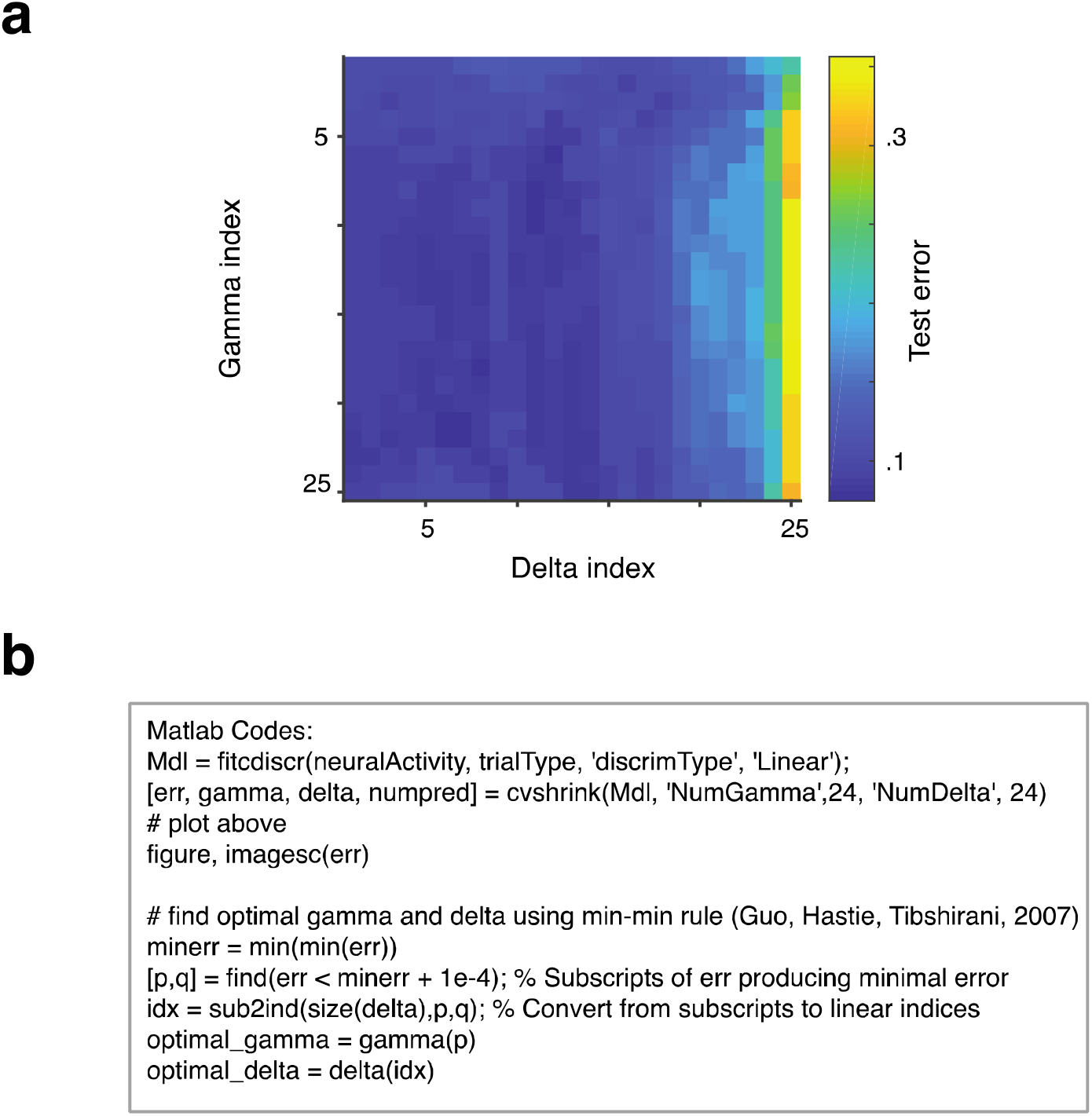
(Supporting figure for Methods) Example of regularization procedure on decoders. **a.** Validation error on Gamma-Delta space (see Methods, Section Decodability) **b.** Matlab code for searching the optimal pair of Gamma-Delta parameters according to the validation errors.

**Supplementary Figure 5.**
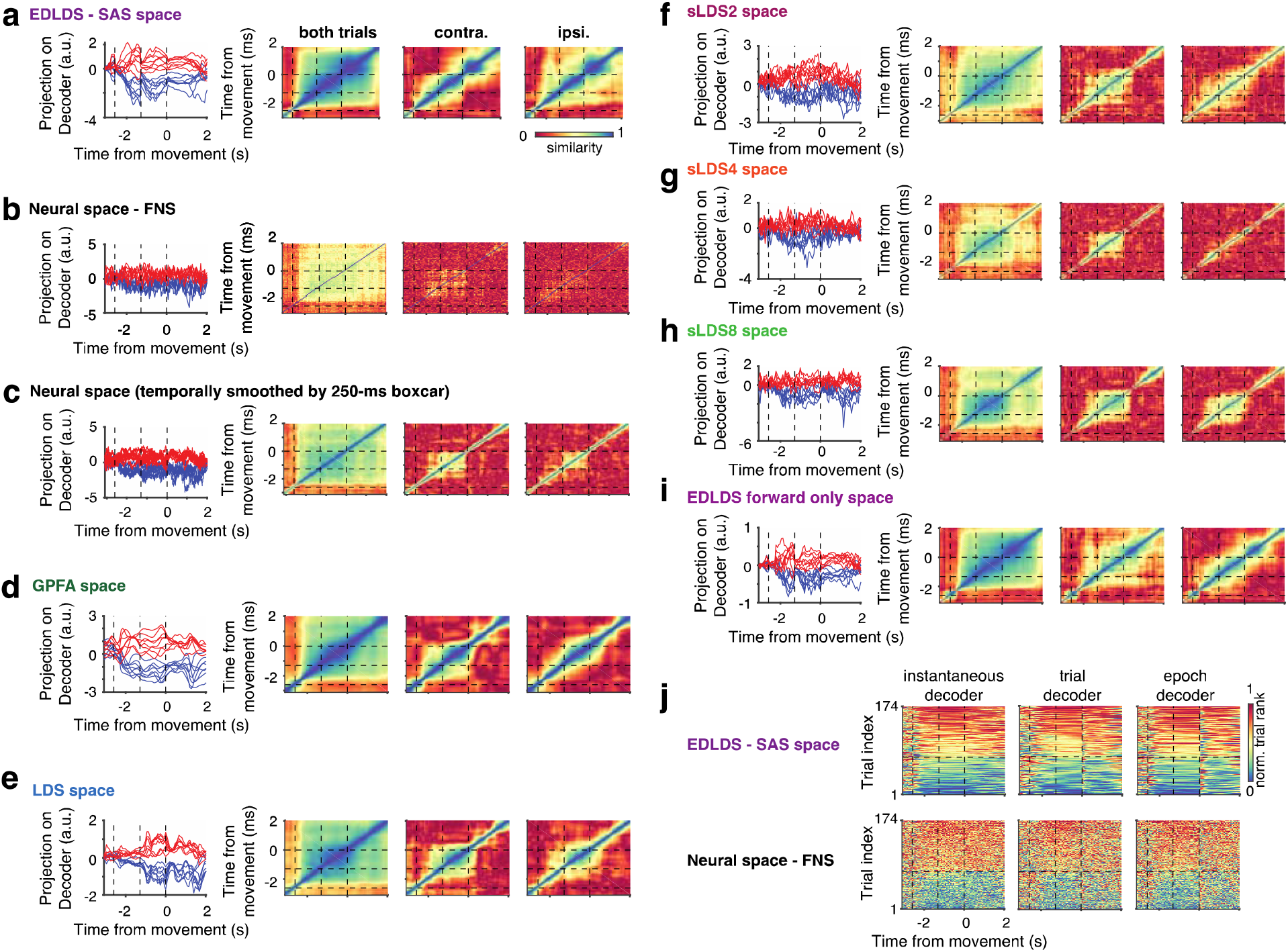
(Supporting figure for Figure 5c-f) Rank correlation of trials in shared activity space. **a. left,** Traces of single trial projections for the two trial types onto trial type decoders based on the EDLDS shared activity space; **right,** Rank correlation between the neural projections onto the instantaneous coding direction at different time points, across both trial types (left), across only contra. trials (middle), and across only ipsi. trials (right). b-i. the same conventions as a. except those based on the full neural space, temporally smoothed full neural space (250 ms boxcared), Gaussian process factor analysis (GPFA) shared activity space, time-invariant linear dynamical systems (LDS) shared activity space, switched linear dynamical system with 2 hidden states (sLDS2), switched linear dynamical system with 4 hidden states (sLDS4), and switched linear dynamical system with 8 hidden states (sLDS8), respectively. **j.** Rank of all trials across time. Data sorted according to average rank in sample-delay epochs. Each plot uses projections on a different decoder: instantaneous decoders (left), decoder based on average activity from sample to response epoch (middle) and a decoder that changes in each epoch and uses the average activity in that epoch (right). Top, EDLDS shared activity space; bottom, full neural space.

**Supplementary Figure 6.**
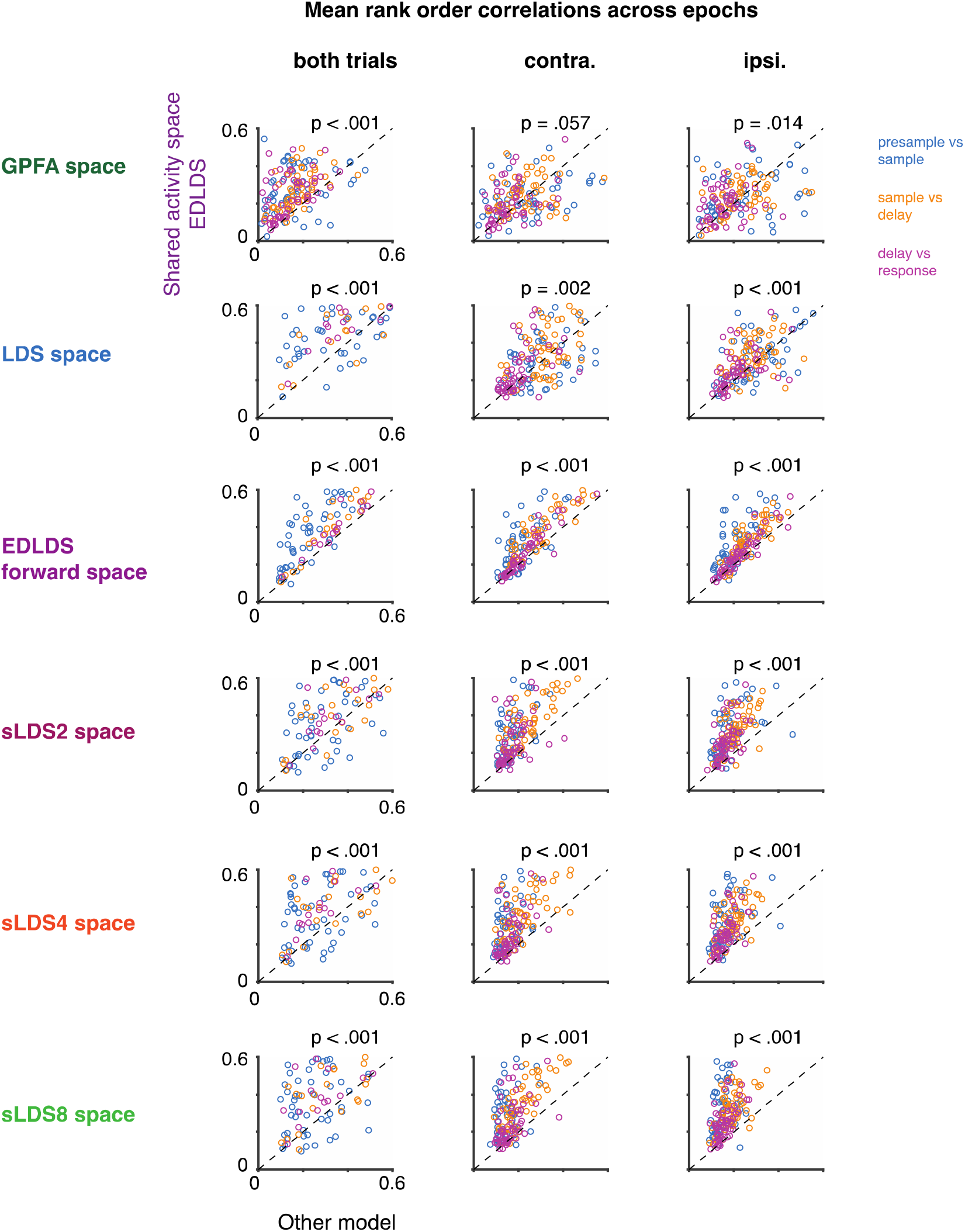
(Supporting figure for Figure4g) Following the same convention as that in Fig. 4g except for other models. P-value is reported using sign-rank test.

**Supplementary Figure 7.**
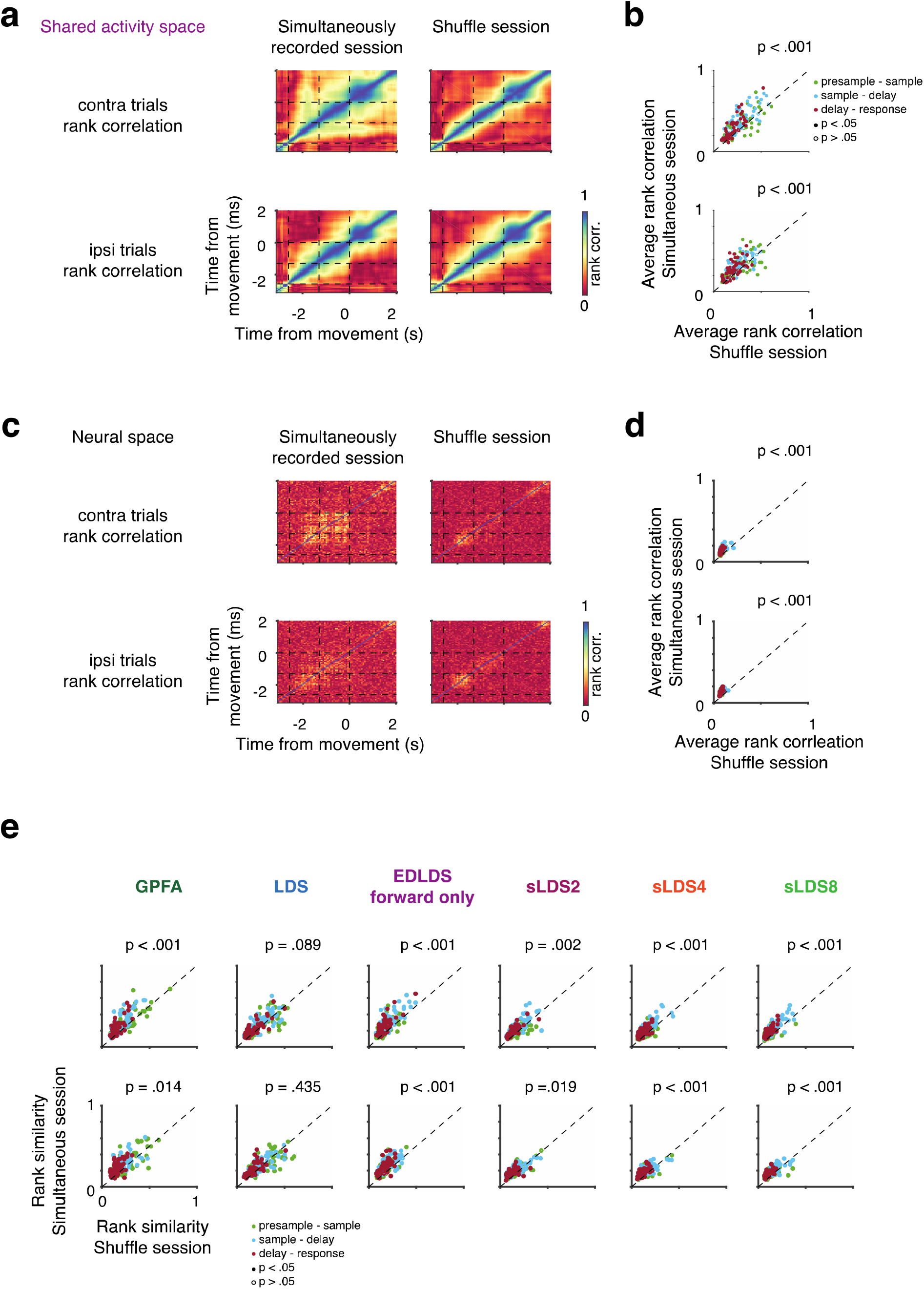
Dynamics of trial identity are disrupted in shuffle trials. **a.** Comparison of rank correlation in simultaneous-recorded trial with shuffle trial session (based on the data from the same session, Session #17). The rank correlation is strong across epochs in simultaneous trials, and is weak in shuffled the trial. Top, contra-only trials; Bottom, ipsi-only trials. **b.** Comparison across all sessions (n = 55). **c-d.** The same convention as Fig a-b except in full neural space. **e.** the same convention as that in **b** except for other models. P-value is reported using sign-rank test.

**Supplementary Figure 8.**
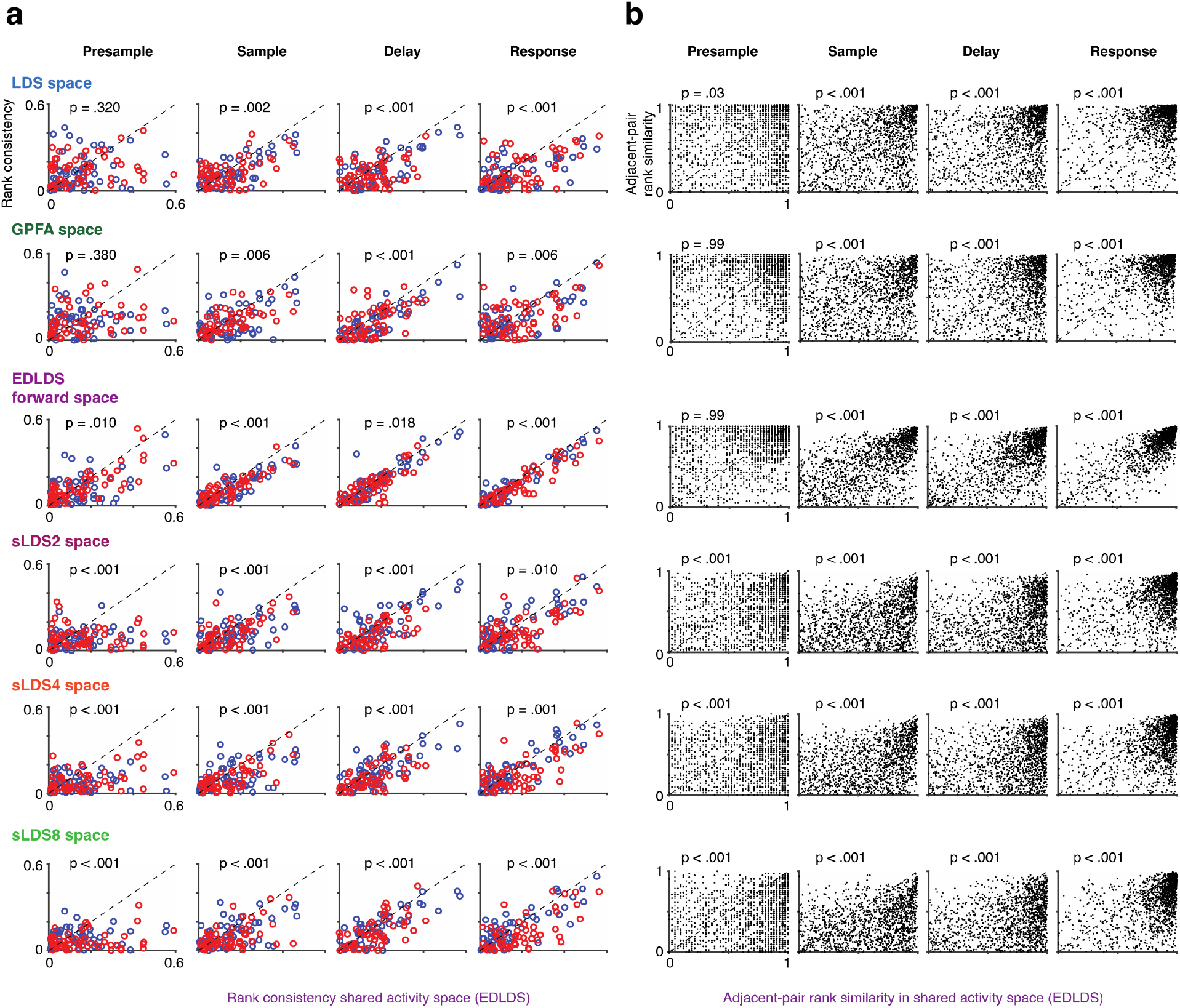
(Supporting figure for Figure 5cd) **a.** Following the same convention as that in Fig. 5c except for other models. P-value is reported using sign-rank test. **b.** Following the same convention as that in Fig. 5d except for other models. P-value is reported using sign-rank test. c. Following the same convention as that in Fig. 5f except for other models. P-value is reported using sign-rank test.

**Supplementary Figure 9.**
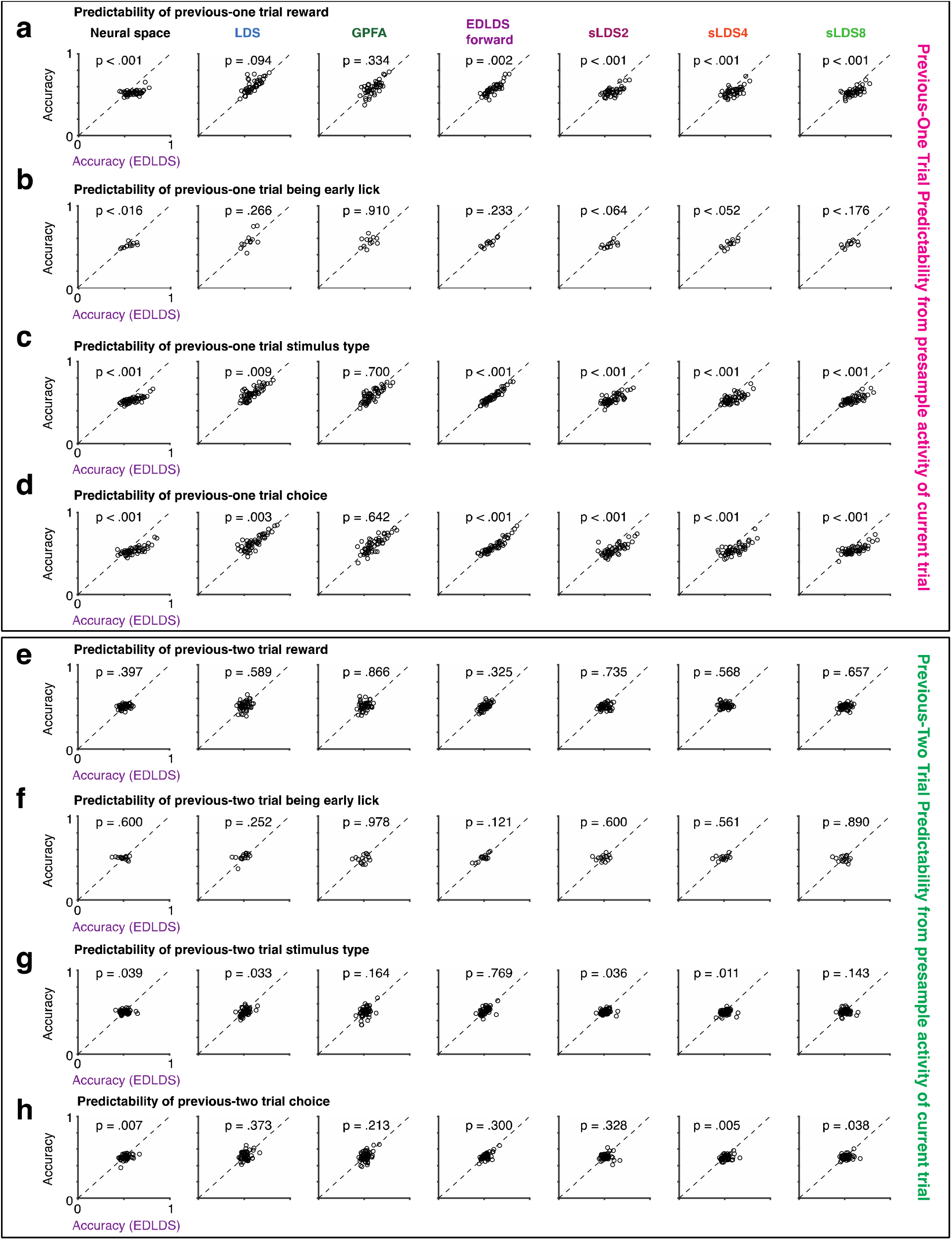
(Supporting figure for Figure 5e-g) **a.** Following the same convention as that in Fig. 5f except for all models. P-value is reported using sign-rank test. **b-d.** Following the same convention as that in **a** except for predictions of previous trial being early lick, previous trial stimulus type and previous trial choice respectively. **e-h.** Following the same convention as that in **a-d** respectively except for predictions of previous-two trial.

**Supplementary Figure 10.**
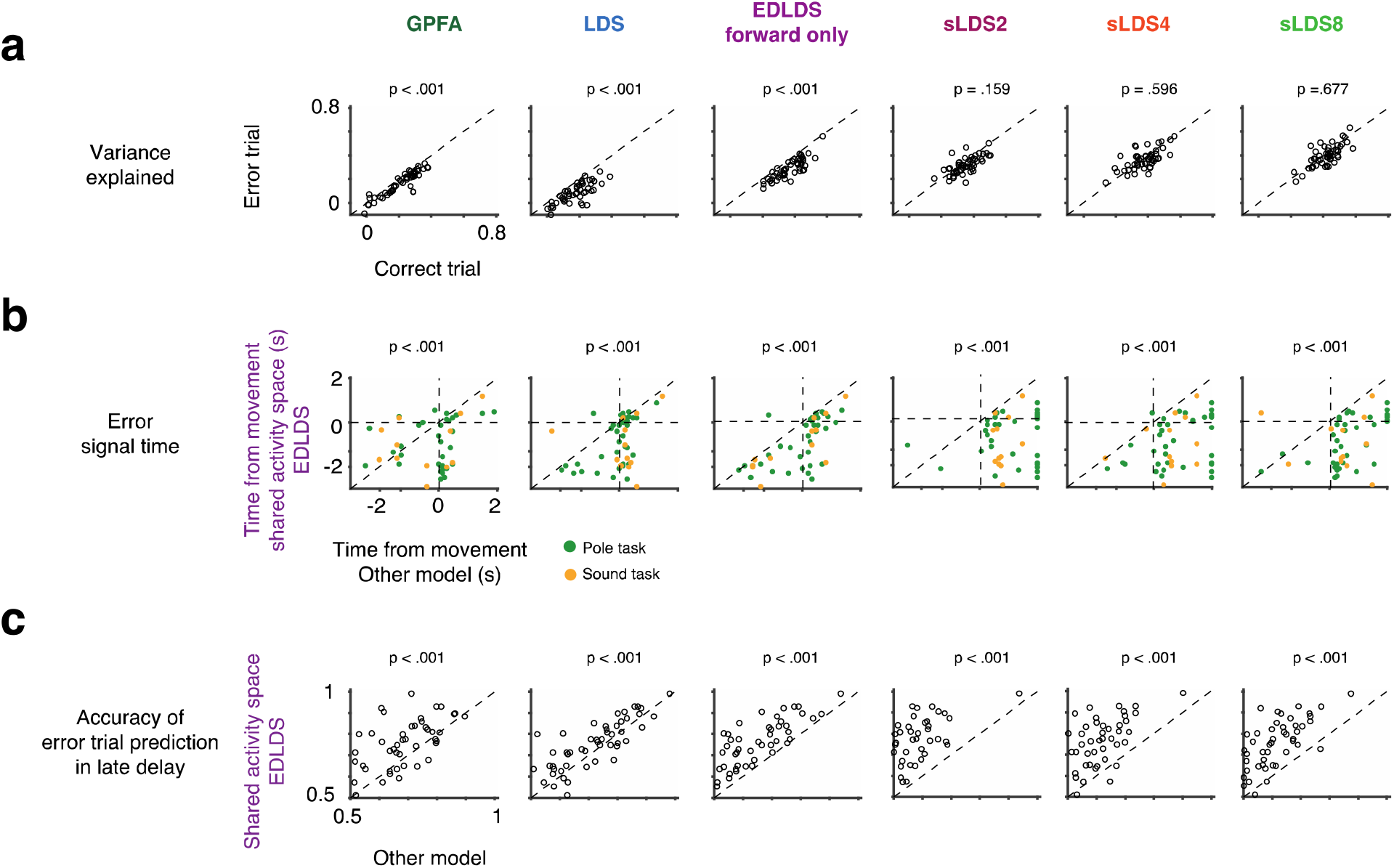
(Supporting figure for Figure 6bde) **a.** Following the same convention as that in Fig. 6b except for other models, **b.** Following the same convention as that in Fig. 6d except for other models, **c.** Following the same convention as that in Fig. 6e except for other models. All p-value is reported using sign-rank test.

**Supplementary Figure 11.**
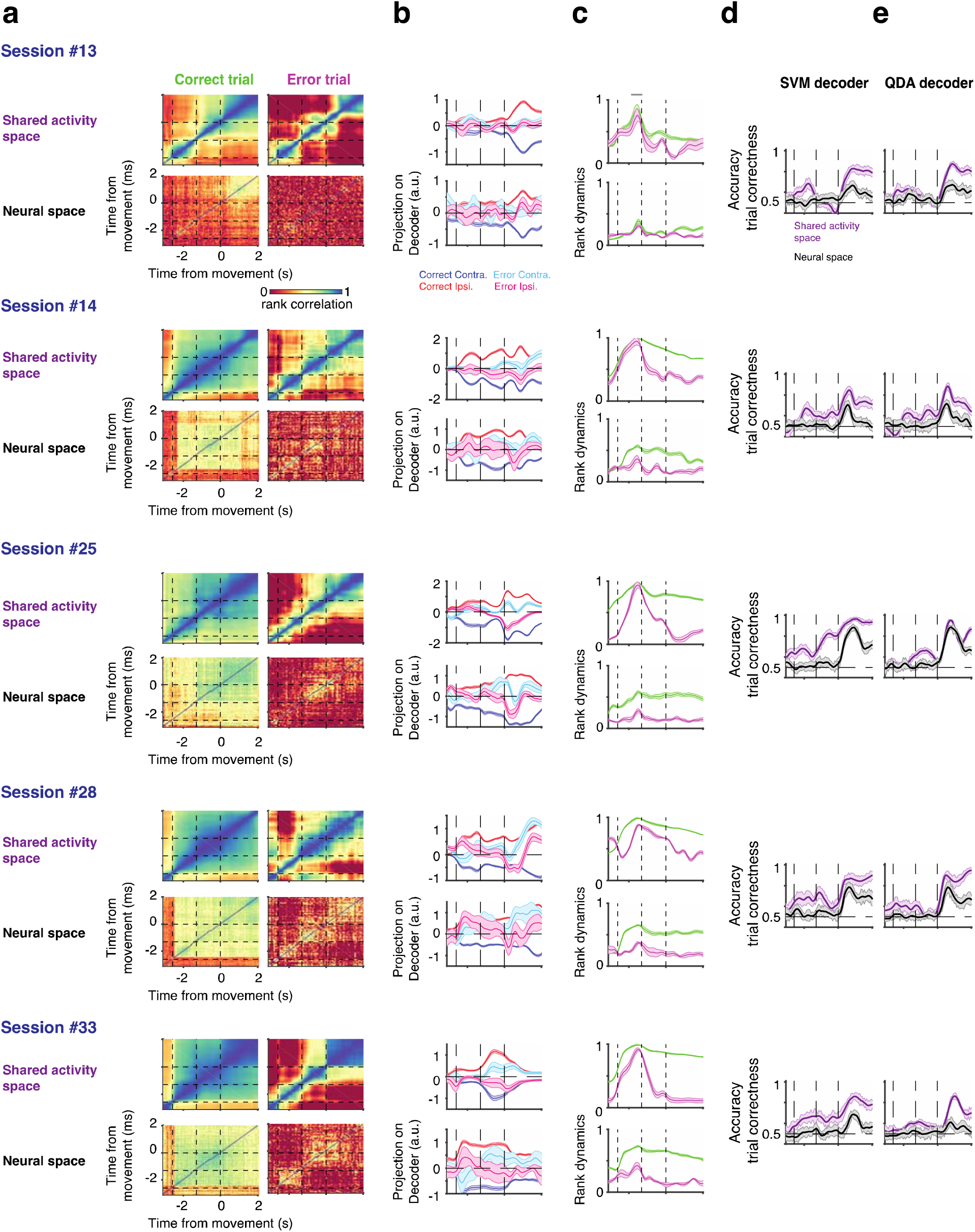
(Supporting figure for Figure 6fghi) Details of neural prediction of correctness on single trials for five more sessions. **a.** the same convention as that in Fig. 6h, except for five more other sessions (Sessions #13, #14, #25, #28, #33). Top, based on neural rank dynamics in EDLDS shared activity space; bottom, based on that in full neural space, **b.** the same convention as that in Fig. 6g, except for five more other sessions. Top, based on neural rank dynamics in EDLDS shared activity space; bottom, based on that in full neural space, **c.** the same convention as that in Fig. 6i, except for five more other sessions. Top, based on neural rank dynamics in EDLDS shared activity space; bottom, based on that in full neural space, **d.** the same convention as that in Fig. 6f, except for five more other sessions, **e.** the same convention as that in **d.,** except using quadratic discriminant analysis to decode error signals.

